# Repurposing the HMG-CoA Reductase Inhibitor Atorvastatin for SRD5A3-CDG

**DOI:** 10.64898/2026.01.18.699766

**Authors:** Hiba Daghar, Seul Kee Byeon, Claudia Maios, Jennifer Poon, James J. Doyle, Wasantha Ranatunga, Éric Samarut, Ethan O. Perlstein, Eva Morava, Akhilesh Pandey, J. Alex Parker, Kristin A. Kantautas

## Abstract

SRD5A3-CDG is a rare autosomal recessive congenital disorder of glycosylation characterized by multisystemic dysfunction, including neurological, psychomotor, cognitive, and visual impairments. Approximately 60 cases have been reported, with treatment limited to symptomatic management. SRD5A3 encodes a polyprenal reductase enzyme essential for synthesizing dolichol, a lipid carrier of the oligosaccharide precursor in N-glycosylation. To address the lack of effective treatments and disease models suitable for high-throughput screening, we developed the first *C. elegans* model of SRD5A3-CDG, harboring the homozygous W19X nonsense mutation commonly observed in patients. This model recapitulates disease-relevant phenotypes, including developmental delays, neurological dysfunction, and mevalonate pathway dysregulation. Using this model, we conducted a high-throughput motility-based drug repurposing screen and identified atorvastatin, an FDA-approved HMG-CoA reductase inhibitor, as a repurposing candidate. Atorvastatin rescued disease-relevant phenotypes in the worm model and restored polyprenol-to-dolichol ratios in patient fibroblasts. These findings highlight atorvastatin as a promising drug repurposing candidate for SRD5A3-CDG.

## Introduction

SRD5A3-CDG is a rare autosomal recessive congenital disorder of glycosylation (CDG), one of over 200 identified CDG subtypes^1,2^. The clinical presentation of SRD5A3-CDG is characterized by a broad spectrum of multisystemic dysfunctions, including neurological, psychomotor, cognitive, and ophthalmological impairments^2,3^. Additional manifestations may include dermal, skeletal, and gastrointestinal abnormalities, contributing to the phenotypic heterogeneity of the disorder. To date, approximately 60 cases have been reported in the literature, with treatment options limited to symptomatic management^4–13^.

SRD5A3-CDG is caused by pathogenic variants in the *SRD5A3* gene, which encodes a polyprenal reductase enzyme involved in dolichol biosynthesis^14–17^. Dolichol is required for the biosynthesis of lipid-linked oligosaccharide precursors in protein N-glycosylation^14,18^ **(Figure S1**) and is also a sugar donor in N-glycosylation and other glycosylation pathways, including O-linked glycosylation, C-mannosylation, and GPI anchor biosynthesis. Consequently, defects in dolichol biosynthesis, as observed in SRD5A3-CDG and other CDG subtypes involving impaired dolichol metabolism, have widespread effects on cellular function^14,17^.

SRD5A3 was initially identified as a polyprenol reductase enzyme, catalyzing the single-step conversion of polyprenol to dolichol^14,15,17^. However, a new study demonstrated that the reduction of polyprenol to dolichol is a multi-step enzymatic process with polyprenol sequentially converted to polyprenal, dolichal, and then to dolichol. SRD5A3 has now been redefined as a polyprenal reductase, catalyzing the reduction of polyprenal to dolichal^17^.

Mutations in *SRD5A3* lead to a dolichol deficiency and an accumulation of polyprenal and polyprenol^17^. Polyprenol can compete with dolichol as a phosphorylated oligosaccharide acceptor, resulting in inefficient glycosylation and the transfer of immature glycans on to proteins^17^.

The development of effective treatments for SRD5A3-CDG has been hindered by the lack of disease models, particularly those amenable to high-throughput screening. To address this, we generated and characterized the first *C. elegans* model of SRD5A3-CDG which harbours the corresponding homozygous W19X mutation that is recurrent in patients. This model recapitulates disease-relevant phenotypes of SRD5A3-CDG, including developmental delays, neurological dysfunction, movement disorders and mevalonate pathway dysregulation, which is essential for isoprenoid and dolichol synthesis^15^.

We conducted a high-throughput motility-based drug repurposing screen using the SRD5A3-CDG worm model. Our analysis of structure-activity relationships of screen hits identified several compounds, including atorvastatin, that share a mevalonate pharmacophore, indicating their potential as HMG-CoA reductase (HMGR) inhibitors. HMGR catalyzes the rate-limiting step in the mevalonate pathway, converting HMG-CoA to mevalonate. Atorvastatin is an FDA-approved statin medication widely used to lower LDL cholesterol levels and reduce the risk of cardiovascular diseases. We demonstrate that atorvastatin rescues disease-relevant phenotypes in the SRD5A3-CDG worm model and restores polyprenol-to-dolichol ratios in patient fibroblasts. These findings highlight atorvastatin as a promising drug repurposing candidate for the potential treatment of SRD5A3-CDG.

## Results

### Negative health phenotypes in C. elegans srdf-3/SRD5A3 mutants

The *C. elegans srdf-3* gene is orthologous to human *SRD5A3,* with the nematode protein sharing 27% identity and 86% similarity to the human protein^19,20^. Both proteins share the enzymatic C-terminal 3-oxo-5-alpha-steroid 4-dehydrogenase domain, suggesting functional conservation^21^. A study in yeast and human cells demonstrated that the conversion of polyprenol to dolichol is a multistep process, with SRD5A3/Dfg10, catalyzing the conversion of polyprenal to dolichal^17^ **(Figure 1A)**. In light of these recent findings, SRD5A3’s role in this pathway has not yet been directly examined in *C. elegans*, but evolutionary conservation of this pathway and enzymatic domain, suggests that its function is conserved.

**Figure 1.**
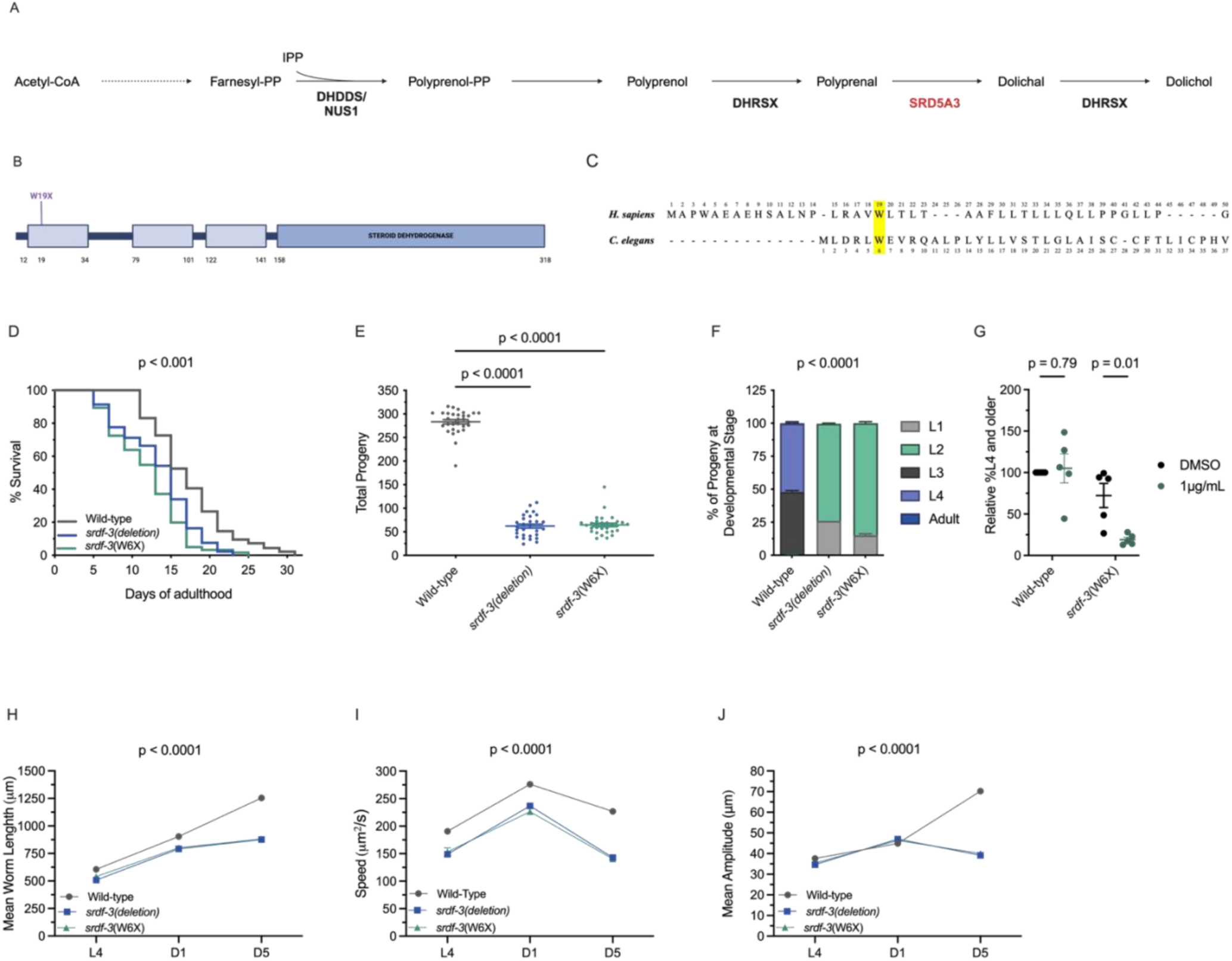
Generation and phenotypic characterization of a SRD5A3-CDG mutation in *C. elegans*. **(A)** Diagram illustrating the function of SRD5A3 in the dolichol biosynthesis pathway, where it catalyzes the conversion of polyprenal to dolichal, a key step in protein glycosylation. **(B)** Location of the patient-specific p.Trp19Ter (W19X) mutation in the first transmembrane domain of human SRD5A3 protein. Boxes represent predicted proteins domains: three transmembrane domains and a steroid dehydrogenase domain. **(C)** Protein alignment showing evolutionary conservation of affected residue (Trp 6) in *C. elegans srdf-3* ortholog (highlighted in yellow). Alignment performed using MARRVEL. **(D)** W6X and *srdf-3*(deletion) mutants have a shorter lifespan relative to wild-type N2 animals. *n* = 120 worms per experiment, *N* = 3 independent experiments. Log-rank (Mantel-Cox) test, two-sided. **(E)** *srdf-3* mutant worms produce significantly fewer offspring than wild-type animals. *n* ≈ 30 worms per experiment, *N* = 3 independent experiments. One-way ANOVA with Bonferonni *post hoc* test. Data are presented as mean ± SEM. **(F)** *srdf-3* mutants exhibit a significant developmental delay compared to wild-type. *n* ≈100 worms per experiment*, N* = 3 independent experiments. Two-way ANOVA with Dunnett *post hoc* test. Data are presented as mean ± SEM. **(G)** Relative percentage of worms reaching L4 and older after treatment with tunicamycin (1µg/mL) or DMSO control. Data are normalized to the respective DMSO conditions for WT and W6X mutant worms. The development of W6X mutant worms is exacerbated by tunicamycin treatment compared to the DMSO control. *n* ≈ 100-300 worms per experiment, *N* = 3 independent experiments. Two-sided unpaired t-test. Data are presented as mean ± SEM. **(H-J)** Mean worm length, speed and crawling amplitude are significantly reduced in *srdf-3* mutants throughout development compared to the wild-type control. The same worms were used across L4, day 1 and day 5 stages. *n* ≈ 106 worms per experiment, *N* = 3 independent experiments. Two-way ANOVA with Tukey *post hoc test.* Data are presented as mean ± SEM.

The most common pathogenic variant in SRD5A3-CDG, p.Trp19Ter (W19X), is a nonsense mutations that results in a loss of protein function^7^. Using CRISPR/Cas9 gene editing, we generated a homozygous W6X *srdf-3* mutant corresponding to human W19X **(Figure 1B, C)**. Additionally, we generated a mutant with a homozygous 1499 bp deletion, resulting in a complete knockout of *srdf-3*, referred to from here on as *srdf-3*(deletion).

The *srdf-3* mutants appear superficially normal; however, they exhibit reduced lifespans compared to wild-type worms **(Figure 1D)**. Progeny production, an indicator of general and reproductive health^22,23^, is also reduced in the *srdf-3*(deletion) and W6X mutants (**Figure 1E)**. Assays measuring larval development rates^24^ revealed a significant developmental delay in both *srdf-3* mutants compared to wild-type controls **(Figure 1F)**.

Since *SRD5A3* mutations disrupt N-glycan biosynthesis and induces ER stress^25^, we investigated whether W6X mutants display related phenotypes. Glycosylation-defective mutants are often hypersensitive to the ER stress inducer tunicamycin, which inhibits glycosylation by promoting protein misfolding and affects post embryonic development in *C. elegans*^16,26–28^. Tunicamycin treatment exacerbated developmental delays in W6X worms, while the development of wild-type worms was not significantly affected **(Figure 1G)**. These findings indicate that the *W6X* mutant is more sensitive to ER stress.

Other health indicators include general morphology and motility. Automated phenotyping measurements of mean worm length at the L4 larval stage, adult Day 1 and Day 5, revealed that *srdf-3*(deletion) and W6X mutants were significantly shorter than wild-type worms throughout development **(Figure 1H, S2)**. For motility, we measured speed **(Figure 1I, S2)** and the mean amplitude of body movements **(Figure 1J, S2)**, both of which were reduced in *srdf-3*(deletion) and W6X mutants compared to wild-type controls during adult stages. The similar phenotypes of the point mutation and deletion suggests that W6X represents a complete loss of *srdf-3* function. Therefore, we proceeded with further phenotypic characterization of the W6X patient-relevant mutant strain.

### Impaired movement and neuronal function in W6X mutants

We assessed W6X worm movement on liquid and solid media to examine progressive motility defects. Swimming assays showed significantly decreased swimming behaviour (thrashing) in W6X mutants on day 1 of adulthood compared to wild-type worms **(Figure 2A)**. On solid media, W6X mutants exhibited a progressive decline in motility, eventually leading to paralysis with age, in contrast to wild-type worms **(Figure 2B)**. This decrease in motility is analogous to the progressive movement deficits observed in SRD5A3-CDG patients.

**Figure 2.**
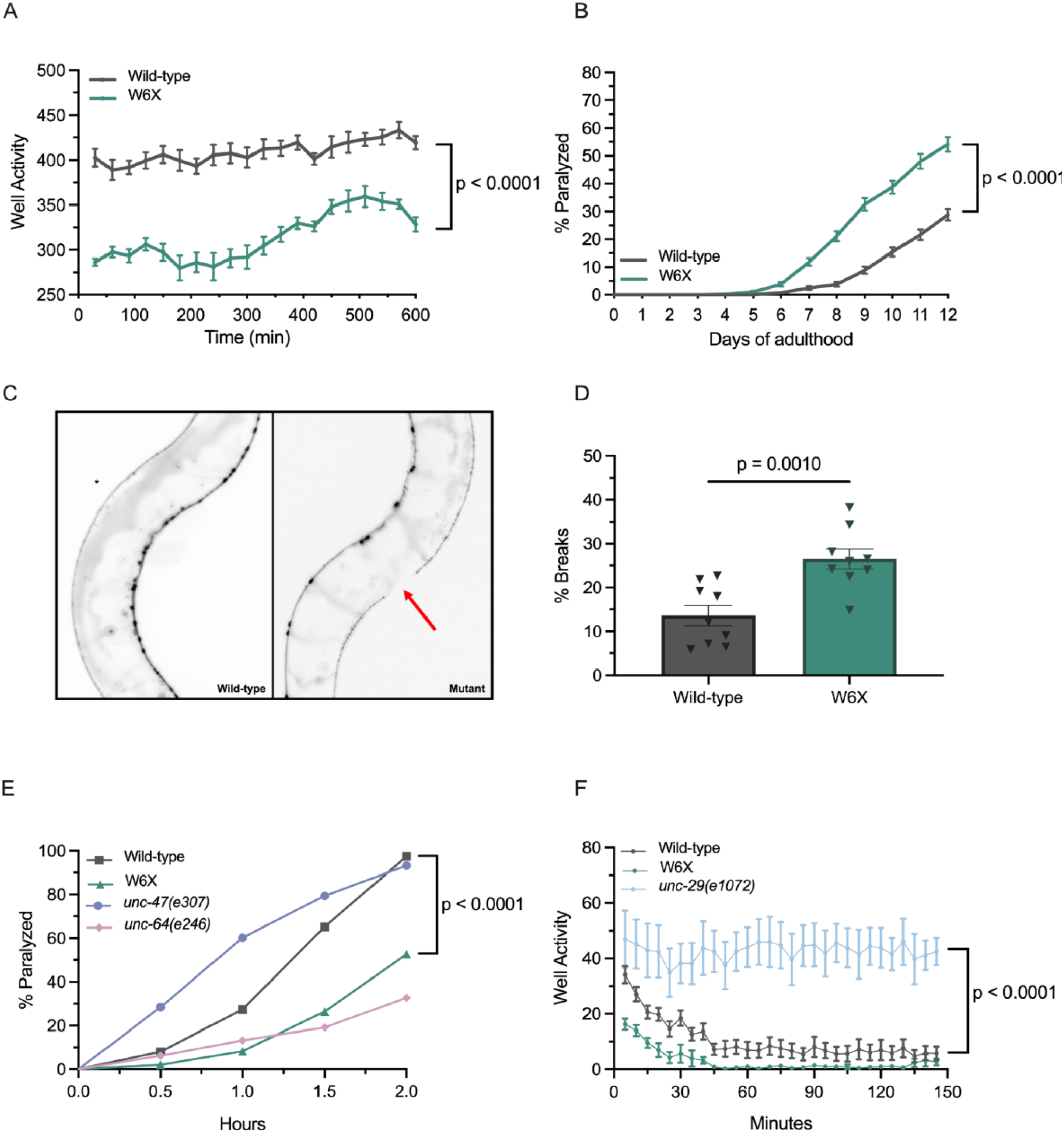
Impaired movement and neuronal function in W6X worms. **(A)** Swimming behavior is significantly impaired in W6X in the early adulthood stage (day 1) *n =*150-210 worms per experiment, N = 3 independent experiments. Two-way ANOVA with Bonferroni *post hoc* test. Data are presented as mean ± SEM. **(B)** W6X mutants cultured on solid media show motility defects leading to paralysis at a higher rate compared to wild type controls. *n* ≈ 120 worms per experiment, *N* = 3 independent experiments. Log-rank Mantel-Cox test, two-sided. Data are presented as mean ± SEM. **(C)** Representative image of a break or gap (red arrow) along a GABAergic motor neuron process in a W6X mutant. (**D**) W6X mutants have a greater frequency of breaks/gaps along neuronal processes compared to wild-type *unc-47::*mCherry controls at day 3 of adulthood. *n* = 30 worms per group, *N* = 3 independent experiments. Two-tailed unpaired t-test. Data are presented as mean ± SEM. **(E)** W6X mutants show significantly less paralysis upon aldicarb treatment compared to wild-type and hypersensitive *unc-47(e307)* mutants, but exhibit similar paralysis levels to resistant *unc-64(e246)* controls*. n* ≈ 120 worms per experiment, *N =* 3 independent experiments. Log-rank Mantel-Cox test, two-sided. Data are presented as mean ± SEM **(F)** W6X mutants exhibited no significant difference in levamisole-induced paralysis compared to wild-type animals, while both showed significantly decreased motility compared to the levamisole-resistant *unc-29(e1072)* control. *n* ≈ 30 worms per experiment, *N* = 3 independent experiments. Two-way ANOVA with Tukey *post hoc* test. Data are presented as mean ± SEM.

Given that SRD5A3-CDG is characterized by neurological symptoms, we investigated neuronal phenotypes in W6X mutants. Since impaired motility can be linked to defective neuronal function or morphology, we used a transgenic reporter strain expressing fluorescent mCherry proteins in GABAergic motor neurons. We observed a significant increase in dorsal cord breaks/gaps in W6X mutants at day 3 of adulthood **(Figure 2C, D)**, suggesting progressive neuronal defects consistent with their paralysis phenotype^29,30^.

To further assess neurological dysfunction at the neuromuscular junction (NMJ), worms were exposed to the acetylcholinesterase inhibitor, aldicarb. Aldicarb causes acetylcholine accumulation at the NMJ, resulting in muscular hypercontraction and rapid paralysis ^31^ (34). W6X mutants exhibited reduced paralysis compared to wild-type worms and hypersensitive *unc-47(e307)* mutants, and resembled aldicarb-resistant *unc-64(e246)* controls **(Figure 2E)**. The aldicarb resistance in W6X suggests synaptic dysfunction, as aldicarb resistance is typically associated with impaired synaptic neurotransmitter release (increased GABA release or reduced acetylcholine release).

To assess whether the aldicarb resistance in W6X mutants is due to presynaptic or postsynaptic dysfunction, we exposed mutants to levamisole, a postsynaptic acetylcholine receptor agonist that induces time-dependent paralysis in wild-type worms^32^. W6X mutants showed no significant difference in levamisole-induced paralysis compared to wild-type worms, while both wild-type and W6X mutant worms displayed reduced motility relative to the levamisole-resistant *unc-29(e1072)* control **(Figure 2F)**. These results suggests that postsynaptic acetylcholine receptors are functional in W6X mutants, supporting the conclusion that their resistance to aldicarb is likely due to presynaptic neuronal defects. These findings are consistent with the observed increase in presynaptic breaks in mutant worms, which may contribute to their motor dysfunction.

### Metabolic dysfunction in W6X mutants

SRD5A3 is involved in the multi-step conversion of polyprenol to dolichol by reducing polyprenal to dolichal. Thus, the ratio of polyprenols-to-dolichols serve as a key indicator of SRD5A3 function. Impaired SRD5A3 activity disrupts this conversion, leading to the accumulation of polyprenol species that compete with dolichol as oligosaccharide acceptors, resulting in inefficient glycosylation and hypoglycosylation^17^.

Previous studies have reported abnormal polyprenol-to-dolichol ratios in SRD5A3 mutant yeast and cellular models, highlighting the imbalance of these metabolites in the context of SRD5A3-CDG^15,17,33^. Furthermore, the accumulation of polyprenols and related intermediates may trigger metabolic rewiring of the mevalonate pathway, a phenomenon observed in other CDG types^34,35^. In *C. elegans*, the non-cholesterol branch of the mevalonate pathway, which generates dolichol, coenzyme Q, and isoprenylated proteins, is conserved, although it lacks the cholesterol biosynthesis branch that is present in mammals^36^.

To assess mevalonate pathway dysregulation in SRD5A3 deficiency, we quantified metabolites in W6X mutants and controls by LC-MS/MS. W6X worms showed significantly increased polyprenols, decreased dolichols, and an elevated polyprenol-to-dolichol ratio, consistent with impaired conversion of polyprenols to dolichols **(Figure 3A-C)**. Several upstream metabolites, including acetoacetyl-CoA, mevalonate and mevalonate-phosphate, were significantly dysregulated in mutant worms **(Figure 3D-F)**. These findings indicate that the impaired conversion of polyprenol to dolichol leads to metabolic rewiring of the mevalonate pathway in worms.

**Figure 3.**
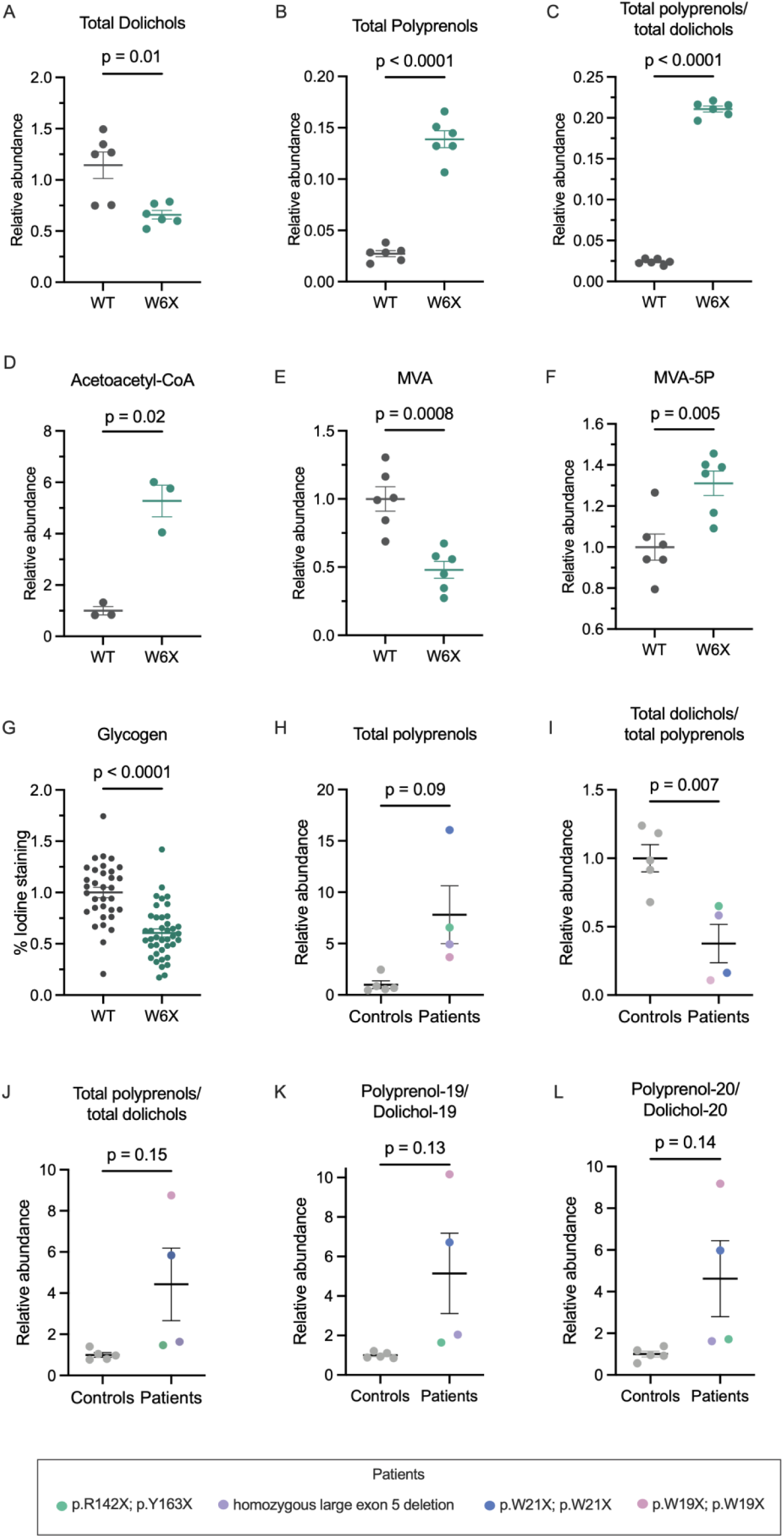
Metabolic dysregulation in SRD5A3-CDG worms and patient fibroblasts. LC-MS/MS analysis revealed **(A)** significantly decreased dolichols, **(B)** increased polyprenols, **(C)** an elevated polyprenols:dolichols ratio and **(D-F)** alterations in acetoacetyl-CoA, mevalonate (MVA), and mevalonate-phosphate (MVA-5P) in W6X mutant worms relative to wild-type (WT)**. a-f:** *n* = 3 technical replicates, *N* = 2 independent experiments. Two-sided unpaired t-test, Welch’s correction. Data are presented as mean ± SEM **(G)** Glycogen abundance is significantly reduced in W6X mutants compared to wild-type worms by iodine staining. *n* ≈ 10 worms per experiment, *N* = 3 independent experiments. Two-sided unpaired t-test. Data are presented as mean ± SEM. In SRD5A3-CDG fibroblasts, LC-MS/MS quantified (**H**) total polyprenols, (**I**) total dolichols:polyprenols ratio, and (**J**) reciprocal ratios and species-specific polyprenol:dolichol ratios with (**K**) 19 and (**L**) 20 isoprene units. **g-i**: *n* = 1 sample per cell line, *N* = 1 experiment. Two-sided unpaired t-test, Welch’s Correction. Data are presented as mean ± SEM.

Increasing evidence suggests that CDGs and glycogen storage diseases (GSDs) share overlapping symptoms^37–40^ with glycosylation defects observed in some GSDs and glycogen metabolism defects reported in CDGs^41,42^. Several glycosylation genes are also involved in glycogen metabolism^43^. Since glycogen metabolism is well-conserved between *C. elegans* and humans^44^, we quantified glycogen in W6X mutant worms and found significant lower glycogen abundance in mutants compared to wild-type worms **(Figure 3G)**, suggesting potential disruptions in glycogen metabolism.

### Metabolic dysregulation in SRD5A3-CDG patient fibroblasts

Dolichols and polyprenols with 18-21 isoprene units were measured from SRD5A3-CDG patient fibroblasts using LC-MS/MS and compared to age- and sex-matched unaffected control fibroblasts (**Supplementary Figure 4A-H and Supplementary Table 1)**. Total levels of dolichols and polyprenols were calculated by summing the abundance of individual species. As expected, total polyprenols directionally higher in the patient cells as compared to controls (an average of 8-fold) although the difference was not statistically significant (p=0.09) (**Figure 3H**). The ratio of total dolichols to total polyprenols was significantly decreased (p<0.05) in the patients (**Figure 3I**), while its reverse ratio of total polyprenols to total dolichols was directionally higher in patients by ∼4-fold with p=0.15 (**Figure 3J)**. Additionally, the ratios of individual polyprenols to dolichols with identical isoprene units, such as polyprenol-19 to dolichol-19 and polyprenol-20 to dolichol-20, trended higher by an average of 3- to 5-fold in patients, although the difference was not statistically significant. (**Figure 3K-L and Supplementary Figure 4I-J**).

### Drug repurposing screen in W6X worms identifies HMG-CoA reductase as a potential therapeutic target

The liquid culture motility assay provided an automated, phenotypic readout for high-throughput drug screening in W6X worms. We screened a library of ∼ 4,000 compounds—including FDA-approved drugs, investigational drugs, bioactives, tool compounds, and natural products—in singlicate to identify compounds that rescue W6X worm motility. From the primary screen, 153 compounds improving motility were selected for a secondary motility screen, conducted in triplicate. Of these, 75 compounds were identified as rescuers, with Z-scores greater than 0 (**Figure 4A**).

**Figure 4.**
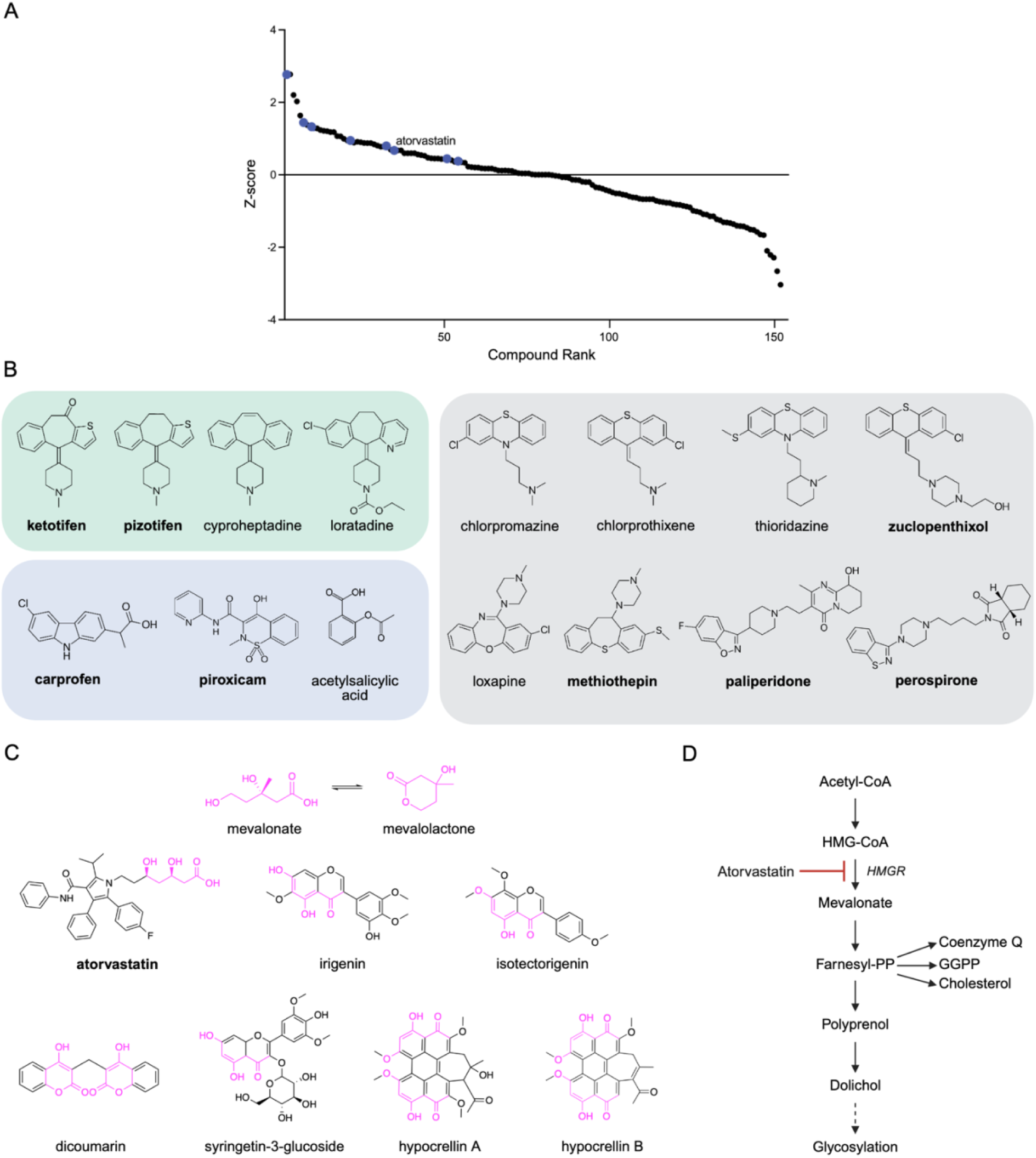
Drug repurposing candidates discovered in a W6X worm drug screen. **(A)** 153 repurposing candidates from the primary screen were advanced to the secondary screen. In both screens, compounds were assessed for their ability to improve W6X worm motility. Compounds from the secondary screen are rank-ordered by Z-score. Blue circles represent the 10 compounds selected for further validation. Atorvastatin is highlighted (Z-score = 0.6745). (**B**) Compound classes of selected repurposing candidates from the secondary screen that rescued worm motility are highlighted. Compounds are categorized as antihistamines (green box), nonsteroidal anti-inflammatory drugs (NSAIDs; blue box), and antipsychotics (grey box). (**C**) The conversion of HMG-CoA to mevalonate is the rate-limiting step in the mevalonate pathway. Mevalonate can exist in both its open-chain form (mevalonate) and its cyclic form (mevalolactone). Structures of rescue compounds from the secondary screen that contain the mevalonate or mevalolactone pharmacophore (highlighted in magenta) are shown. (**D**) A simplified schematic of the mevalonate pathway and dolichol biosynthesis. Atorvastatin, a competitive inhibitor of HMG-CoA reductase, reduces flux through the mevalonate pathway. Bolded compound names in **B-C** indicate compounds selected for further validation: atorvastatin, ketotifen, pizotifen, zuclopenthixol, methiothepin, paliperidone, perospirone, carprofen and piroxicam. Z-scores of and the percent average movement activity of compounds in B-C are reported in Table S2.

Several hit classes emerged among the identified rescue compounds, including antihistamines, antipsychotics, non-steroidal anti-inflammatory drugs (NSAIDs) (**Figure 4B, Table S2**). We also observed several compounds which share a complete or partial mevalonate pharmacophore, including plant-based polyphenols (irigenin, isotectorigenin, dicoumarin, and syringetin-3-glucose), fungal-derived perylenequinones (hypocrellin A and hypocrellin B) and atorvastatin (**Figure 4C, Table S2**). Atorvastatin showed reproducible motility rescue, with a Z-score of 3.4 in the primary screen and in 0.67 the secondary screen (**Table S2**). The conversion of HMG-CoA to mevalonate by HMGR is the rate-limiting step in the mevalonate pathway and cholesterol biosynthesis. Atorvastatin, an FDA-approved statin, is a competitive HMGR inhibitor that is used to lower blood cholesterol levels^45^ (**Figure 4D**). Plant-based polyphenols, known for their complex polypharmacology, as well as antioxidant and anti-inflammatory properties, have been associated with reduced cholesterol levels and improved cardiovascular health through dietary consumption^46–48^. Molecular docking studies suggest that dietary polyphenols may competitively inhibit HMGR^49,50^. While HMGR inhibition can reduce cholesterol levels by decreasing flux through the mevalonate pathway, it may also lower polyprenol levels. Taken together, the structure-activity relationships of these compounds highlight HMGR as a potential therapeutic target for SRD5A3-CDG.

### Atorvastatin rescues W6X worm phenotypes

SRD5A3 deficiency leads to polyprenol accumulation, which competes with dolichol as a phosphorylated lipid-linked oligosaccharide acceptor, leading to the transfer of immature glycans and hypoglycosylation^17,33,51,52^. We hypothesized that reducing flux through the mevalonate pathway at the rate-limiting step by inhibiting HMGR, could mitigate the effects of polyprenol accumulation when SRD5A3 activity is diminished or absent. Based on this hypothesis and because atorvastatin is an FDA-approved drug with a well-established safety profile in both adult and pediatric populations, we sought first to validate its effect on the progeny phenotype in W6X worms. Eight other repurposing candidates from recurring drug classes—ketotifen, pizotifen, carprofen, piroxicam, zuclopenthixol, methiothepin, paliperidone, and perospirone—were selected for validation based on their approval status, safety profile, route of administration, and potential for pediatric use. Atorvastatin rescued the reduced progeny phenotype at 0.1 µM but had a toxic effect at higher concentrations (50 µM and 100 µM) (**Figure 5A**). Among the other compounds tested, the NSAID piroxicam also rescued the reduced progeny phenotype at 1 µM and 10 µM (**Figure S3**).

**Figure 5.**
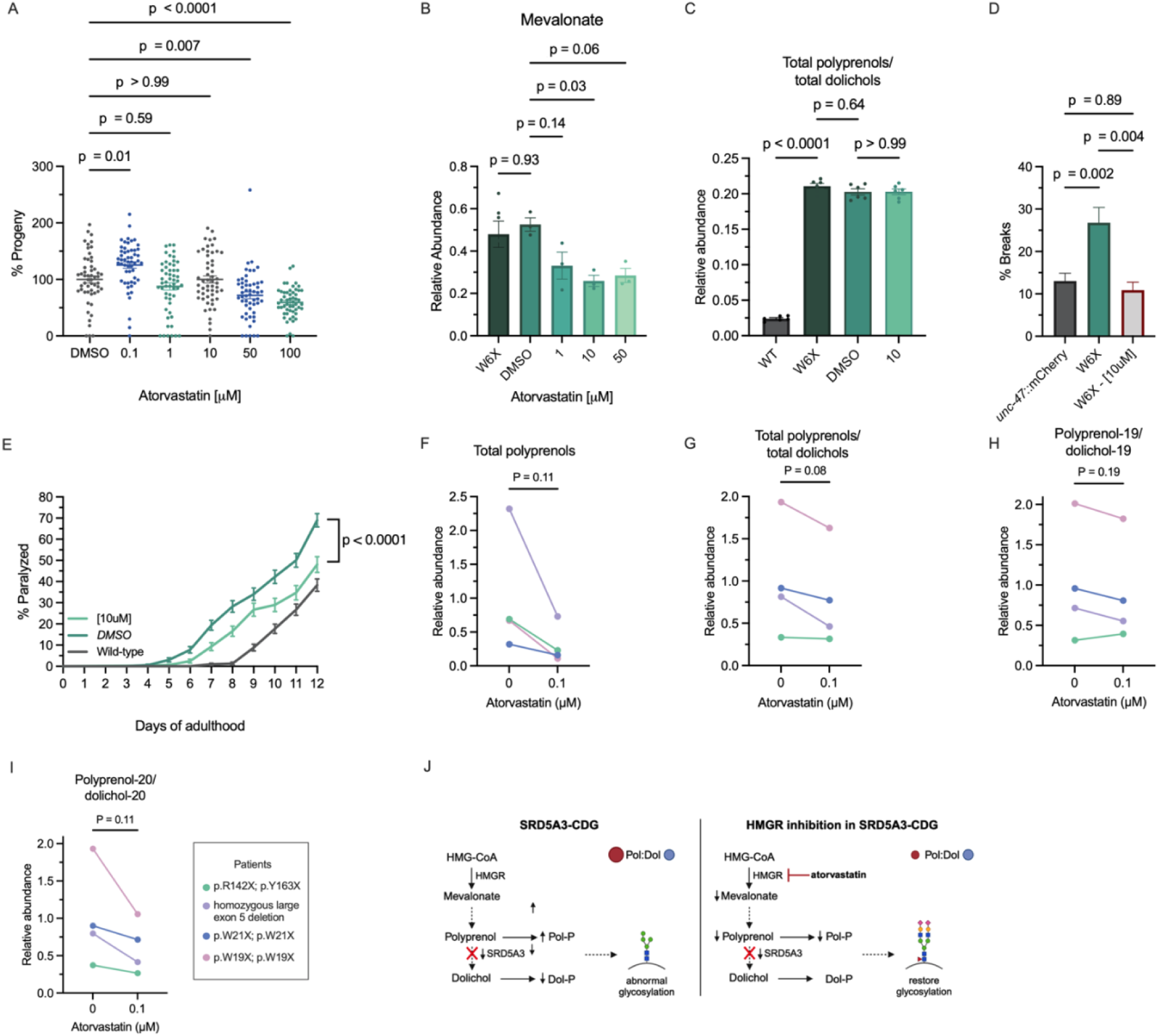
Atorvastatin rescues W6X worm phenotypes and reduces polyprenol:dolichol ratios in SRD5A3-CDG patient fibroblasts. **(A)** Treatment with a low-dose of HMG-reductase inhibitor atorvastatin (0.1µM) rescues the progeny phenotype in W6X mutant worms. *n* = 20 worms per experiment, *N* = 3 independent experiments. Brown-Forsythe one-way ANOVA with Dunnett *post hoc* test. Data are presented as mean ± SEM. **(B)** Atorvastatin (10 µM) treatment decreased mevalonate levels in W6X worms, as quantified by LC–MS/MS (n = 3 technical replicates, N = 1 independent experiment). **(C)** Atorvastatin treatment did not significantly alter the total polyprenol:dolichol ratio in W6X worms, as quantified by LC–MS/MS (n = 3 technical replicates per experiment, N = 2 independent experiments). **b-c**: Brown-Forsythe one-way ANOVA with Dunnett *post hoc* test. Data are presented as mean ± SEM. **(D)** Atorvastatin (10 µM) reduced the number of breaks associated with neuronal morphology defects in W6X mutant worms. *n* ≈ 30 worms per experiment, *N =* 3 independent experiments. Brown-Forsythe one-way ANOVA with Tukey *post hoc* test. Data are presented as mean ± SEM. **(E)** Atorvastatin (10 µM) rescued age-dependent paralysis in W6X mutants compared to the vehicle-treated (DMSO) control. n ≈ 120 worms per experiment, *N =* 3 independent experiments. Log-rank (Mantel-Cox) test, two-sided. Data are presented as mean ± SEM. (**F**) In patient fibroblasts, atorvastatin treatment (0.1 µM) showed a trend towards decreased total polyprenols, (**G**) decreased total polyprenol:dolichol ratio and reduced ratios of individual polyprenol:dolichol species with (**H**) 19 and (**I**) 20 isoprene units. **f-i:** *n* =1 sample per cell line, *N =* 1 experiment. Two-sided paired t-test. Data are presented as mean ± SEM. (**J**) Hypothetical schematic representation of the proposed rescue mechanism of atorvastatin in SRD5A3-CDG. (Left) In SRD5A3-CDG human cells, impaired SRD5A3 function leads to the accumulation of polyprenal and polyprenol, resulting in an increased ratio of polyprenol to dolichol and polyprenol-phosphate (Pol-P) to dolichol-phosphate (Dol-P). Phosphorylated polyprenol competes with dolichol as a lipid carrier for N-glycan assembly, leading to abnormal glycosylation. (Right) Treatment with the HMG-CoA reductase inhibitor atorvastatin, may reduce polyprenol levels, decreases the ratio of polyprenol to dolichol and help restore glycosylation by reducing the competition between Pol-P and Dol-P.

To confirm that atorvastatin’s mechanism of action as an HMGR inhibitor was responsible for the observed phenotypic rescue in worms, we measured mevalonate levels in W6X worms treated with atorvastatin. As expected, mevalonate levels were significantly reduced upon treatment (**Figure 5B**), consistent with inhibition of HMGR. We next examined whether atorvastatin modulated dolichol-pathway metabolites in W6X worms. Treatment with 10 µM atorvastatin did not significantly alter total polyprenols, total dolichols, or the polyprenol to dolichol ratio relative to vehicle-treated W6X controls (**Figure 5C; Figure S9**), indicating that atorvastatin does not restore these metabolites in the worm model. Given that atorvastatin has an on-pathway mechanism of action and successfully rescued the progeny phenotype, we further assessed its effect on additional worm phenotypes. Treatment with 10 µM atorvastatin rescued both the number of neuronal breaks in W6X mutant worms and age-dependent paralysis (**Figure 5C-D**). These results demonstrate that low-dose atorvastatin improves general health and neuronal phenotypes in W6X *C. elegans* mutants, despite having no effect on polyprenol or dolichol levels.

### Atorvastatin rebalances polyprenol-to-dolichol ratios in SRD5A3-CDG patient fibroblasts

Polyprenols and dolichols were measured in patient fibroblasts treated with 0.1 µM atorvastatin for 48 hours and compared to untreated patient fibroblasts. Polyprenols, which were directionally higher in patients compared to unaffected controls, appeared lower following treatment (**Figure 5E)**. Upon treatment, the polyprenol-to-dolichol ratio trended toward rescue as three out of four patient fibroblasts showed 15-40% reduction, although the change was not statistically significant (p=0.08) (**Figure 5F**). The ratios of individual polyprenol-to-dolichol with 19 and 20 isoprene units were examined (**Figure 5G-H**). A subset of patients showed a greater degree of changes in the polyprenol-20 to dolichol-20 ratio, suggesting that the impact of atorvastatin may vary depending on the length of the isoprene units.

## Discussion

In this study, we generated the first *C. elegans* model of SRD5A3-CDG to gain insights into disease mechanisms and discover drug-repurposing candidates for potential therapeutic intervention. We investigated disease-related phenotypes in worms at the motor, cellular, and molecular levels and demonstrated that *srdf-3* worms with the patient-specific loss-of-function W6X mutation, recapitulate key clinical features of SRD5A3-CDG, including developmental delays, neurological dysfunction and metabolic dysregulation of the mevalonate pathway. We also identified decreased glycogen levels as a novel metabolic impairment in *srdf-3* mutant worms. Recent studies indicate a close relationship between glycosylation and glycogen metabolism^40,42,53,54^. The reduced levels of glycogen suggest that glycogen breakdown into glucose may be a compensatory mechanism in SRD5A3-CDG in response to the increasing demand for sugars utilized in N-glycosylation^6^. This metabolic adaption highlights the broader impact of impaired glycosylation on energy metabolism and further supports the notion that metabolic requiring occurs in response to glycosylation defects.

We measured polyprenol and dolichol levels in SRD5A3-CDG patient fibroblasts across various genotypes and polyprenol levels appeared higher by an average of 8-fold in patients to varying extents, while dolichol levels were comparable to healthy controls, consistent with previous reports^17,33^. Surprisingly, fibroblasts with the W21X genotype exhibited polyprenol levels approximately four times higher than those observed in the W19X fibroblast line despite being loss-of-function mutations and located close to exon 1. In agreement with earlier studies, the patient fibroblasts displayed higher polyprenol-to-dolichol ratios^15^. Notably, the patient lines with early truncating mutations showed greater elevations in polyprenol-to-dolichol ratios compared to those with later truncating mutations (p.R142X and p.Y163X) or exon 5 deletions. Interestingly, despite these early termination mutations, dolichol was still present in both patient fibroblasts and worms with loss-of-function mutations, suggesting that dolichol synthesis is not entirely dependent on SRD5A3 activity. Recent glycoproteomic analyses of the same patient fibroblast lines demonstrated widespread abnormalities in N-glycoprotein maturation, confirming glycosylation defects at the cellular level^55^. Future work evaluating whether atorvastatin improves glycosylation in SRD5A3-deficient fibroblasts will be valuable for determining the therapeutic relevance of our findings. The variability in polyprenol and dolichol levels and the fact that dolichol levels remain comparable to or even elevated in some patients may reflect compensatory changes in dolichol biosynthesis. This could involve the activity of enzymes such as DHRSX or the presence of alternative biosynthetic pathways^17,33^.

We conducted an unbiased, motility-based high-throughput screen in W6X worms and identified several prominent hit classes, including antihistamines, antipsychotics, NSAIDs, and polyphenols. These hit classes have also emerged in drug repurposing screens for other CDG types in both worm and fly models. The identification of structurally and mechanistically diverse rescuers suggests that SRD5A3 deficiency perturbs multiple downstream cellular processes, not solely dolichol biosynthesis.

Plant-based polyphenols were the largest hit class in repurposing screens for PMM2-CDG and NGLY1 deficiency^56,57^. The rescue effect of this compound class in these models may be attributed to their antioxidant activity, likely mediated through activation of cytoprotective responses such as the Keap1-Nrf2 pathway or by inhibiting aldose reductase. NSAIDs also emerged as a prominent hit class in drug repurposing screens for NGLY1 deficiency^56^. In our study, we found that piroxicam rescued the progeny phenotype in W6X worms. The mechanism by which anti-inflammatory drugs exert their rescue effect in NGLY1 deficiency, proposed by Iyer et al., 2019 and others, involves the suppression of elevated innate immune responses resulting from impaired mitochondrial function. Mitochondrial dysfunction has been implicated in several CDG types^40,58,59^. Comparative analysis of rescue hit classes across drug repurposing screens for various CDG types, and mitochondrial diseases further underscore the strong relationship between mitochondrial dysfunction and CDG pathology^60^. Investigating mitochondrial function in SRD5A3-CDG, and other CDG types warrants further investigation.

We identified the HMG-CoA reductase inhibitor, atorvastatin, as a promising drug repurposing candidate for SRD5A3-CDG. Our data show a clear reduction in mevalonate levels with atorvastatin treatment in worms, consistent with on-target HMGR inhibition. Although genetic validation of *hmgr-1* could not be performed due to embryonic lethality, prior work establishes HMGR-1 as the physiological target of statins in *C. elegans*, supporting the conclusion that atorvastatin acts through this pathway in our model^61^. Atorvastatin rescued the reduced progeny phenotype, improved motility and reduced neuronal damage in W6X worms, and restored the polyprenol-to-dolichol balance in patient fibroblast lines. However, our results also showed that high doses of atorvastatin had toxic effects, suggesting that optimal therapeutic efficacy requires low-dose statin therapy to reduce flux through the mevalonate pathway without completely shutting it down. This is consistent with the known regulatory dynamics of the pathway, where maintaining some level of activity is crucial for cellular homeostasis. Statins have been successfully used in children with diseases such as hypercholesterolemia and Smith-Lemli-Opitz syndrome^62^, which highlights their potential safety and efficacy in pediatric populations. Given atorvastatin’s approval for use in some pediatric populations, its repurposing for SRD5A3-CDG is promising.

Therapeutic flux modulation within the mevalonate pathway has also been explored in other CDG contexts. In DPM1-CDG fibroblasts, where the synthesis of dolichol-phosphate-mannose is impaired, Haeuptle et al. demonstrated that inhibiting squalene synthase with zaragozic acid redirects flux away from the cholesterol branch of the mevalonate pathway and increases dolichol-phosphate availability, improving LLO maturation and N-glycosylation^63^. Although these findings support the broader principle that adjusting pathway flux can rescue glycosylation defects, the relevance of this strategy to SRD5A3 deficiency remains uncertain, as increasing flux upstream of SRD5A3 could theoretically exacerbate polyprenol accumulation. Statin treatment has previously worsened clinical outcomes in at least one DHDDS-CDG patient, emphasizing that therapeutic manipulation of this pathway is highly gene- and context-dependent^64^. In our study, atorvastatin improved worm phenotypes without altering polyprenol or dolichol levels, while in patient fibroblasts it partially normalized the polyprenol-to-dolichol ratio. This finding is consistent with the polyprenol-phosphate to dolichol-phosphate ratio-based pathomechanism proposed by Wilson et al. (2024)^17^. Together, these findings suggest that carefully titrated modulation of mevalonate pathway flux may offer therapeutic benefit in SRD5A3-CDG, but will require further evaluation in human cellular and clinical settings.

The appearance of several mevalonate-pharmacophore–containing rescuers in the worm screen suggests that modulating flux at HMG-CoA reductase may influence multiple downstream branches of the pathway in SRD5A3 deficiency. The discrepancy between worm and fibroblast metabolite responses to atorvastatin indicates that the consequences of mevalonate-pathway modulation are context-dependent and may reflect differences in how *C. elegans* and human cells utilize downstream isoprenoid-derived products such as prenylated proteins, ubiquinone, and factors involved in mitochondrial maintenance. In worms, phenotypic rescue by atorvastatin may therefore arise through effects on prenylation, mitochondrial function, or neuronal homeostasis rather than through changes in polyprenol or dolichol abundance. Conversely, the polyprenol-to-dolichol ratio improvement observed in patient fibroblasts suggests that atorvastatin can modulate polyprenol-to-dolichol balance in human cells, consistent with the ratio-based model of SRD5A3-CDG pathogenesis and reinforcing its therapeutic potential.

Lastly, increasing the consumption of polyphenol-rich foods or modulating dietary cholesterol intake could serve as alternative therapeutic approaches. Polyphenols may inhibit HMGR activity^49^, while cholesterol modulation could leverage the negative feedback mechanism in cholesterol synthesis regulation. These dietary strategies offer potential complementary methods to fine-tune the balance of the mevalonate pathway and mitigate glycosylation defects in SRD5A3-CDG.

Overall, our findings underscore the utility of *C. elegans* as a model organism for studying glycosylation disorders and identifying drug candidates through high-throughput screening. By investigating numerous disease-related phenotypes in worms and patient cell lines, we provide a comprehensive understanding of SRD5A3-CDG pathophysiology and present atorvastatin as a potential therapeutic option that warrants further exploration in clinical studies.

### Limitations of the study

Since this study was conducted prior to the discovery of multi-step conversion from polyprenol to dolichol, which involves polyprenal and dolichal as intermediates, our targeted LC-MS/MS methods were solely focused on polyprenols and dolichols. As a result, measurements on polyprenals and dolichals are not available. Additional method development is required to investigate any changes in polyprenals and dolichals following the treatment with atorvastatin in worms and patient fibroblasts. Additionally, the full breadth of molecular consequences in this model remains to be characterized, especially to understand the impact on the nervous system, which could explain the vast variety of potential compounds that showed a positive hit from our high throughput experiments. Finally, since *C. elegans* lack the cholesterol biosynthesis pathway branching from the mevalonate pathway present in humans, insights into the effects of atorvastatin on cholesterol regulation in SRD5A3 deficiency and potential feedback mechanisms as a consequence of metabolic rewiring are limited.

## Methods

### Amino acid alignment

BLAST and MARVELL were used for amino acid alignment of SRD5A3 from *Homo Sapiens* (*SRD5A3*; Gene ID: 79644) and *C. elegans* (*srdf-3*; Gene ID: 179438).

### Nematode strains and maintenance

Worms were maintained on standard NGM plates with OP50 *E. coli*. Strains were maintained at 15°C and assays were performed at 20°C. Strains used for the study were N2, CB246(*unc-64(e246))*, CB307(*unc-47(e307))* and CB1072*(unc-29(e1072))*, all provided by the *Caenorhabditis Genetics Center (University of Minnesota)*, which is funded by *NIH Office of Research Infrastructure Programs (P40 OD010440)*. The *srdf-3* deletion mutant (*srdf-3(mod1))* and W6X point mutant (*srdf-3(mod2))* were generated by *SunyBiotech.* IZ629 *ufIs34[Punc-47p::*mCherry] was kindly gifted from Dr. Michael M. Francis (University of Massachusetts, Worcester, MA)^65^.

### Worm genotyping

PCR was used to confirm deletion mutants, while high-resolution melting (HRM) and genome sequencing was used for point mutations. HRM MeltDoctor reagents (Applied Biosystems) and HRM software (Applied Biosystems) were used before verification by Sanger sequencing (Genome Quebec). Primers: HRM-F (CAATTATTCATTTCAGAATGTTAGATCGATTATAA), HRM-R (CTAACAGAT AAAGGGGCAGTGC).PCR-F (CAATTATTCATTTCAGAATGTTAGATCGATTA TAA) and PCR-R (TCCAAGCGTAGATACTAACAGATAA GG). Crosses were performed with IZ629 (*ufIs34[Punc-47p::*mCherry]) to generate *srdf-3*(W6X)*; Punc-47p::*mCherry mutants which were outcrossed five times to establish stable lines.

### Liquid culture motility assay

Synchronized day 1 adult worms were collected, washed with M9 buffer and transferred into 96-well plates in 100 µL M9 buffer (∼50 to 70 worms/well). Motility was recorded for 10 hours with WMicrotracker ONE (*Phylum Tech*). Each condition was tested in triplicate across three independent experimental repeats.

### WormLab analysis

Movement parameters, including worm length, amplitude and speed, were quantified using *MBF Bioscience WormLab* software (2020 version). A 30-second video was recorded for each worm and paramaters were manually tracked. Data were analyzed with GraphPad Prism 9.

### Lifespan analysis

Forty L4-synchronized worms were transferred to NGM plates and transferred to new plates every other day from adult day one until death. Worms were scored as alive, dead, or lost worms. Each condition was tested in triplicate (40 worms/plate) across three independent experimental repeats.

### Development assay

L1-synchronized *C. elegans* were transferred to NGM plates and counted daily by developmental stage until N2 worms reached day 1 of adulthood. Each condition was tested in triplicate (100 worms/experiment) across three independent experimental repeats.

### Progeny assay

Thirty L4-synchronized worms were placed on individual NGM plates. Eggs hatched by day 1 of adulthood were counted as viable progeny. Each condition was tested in triplicate across three independent experimental repeats.

### Paralysis assay

35-40 worms were picked to NGM plates on day 1 of adulthood and scored daily for movement over 12 days. Worms were scored as paralyzed if unresponsive to prodding with a worm pick and dead if they showed no head movement after prodding on the nose and no pharyngeal pumping. Each condition was tested in triplicate across three independent experimental repeats.

### Aldicarb sensitivity assay

40 day 1 adult worms were transferred to 1 mM aldicarb NGM plates. Paralysis was assayed every 30 minutes for 2 hours. Animals were counted as paralyzed if they failed to crawl upon prodding with a platinum wire. Each condition was tested in triplicate across three independent experimental repeats.

### Levamisole sensitivity assay

Synchronized day 1 of adulthood worms were transferred to standard 96-well plates containing 100 µM of levamisole. Motility was recorded for 145 minutes using a WMicrotracker ONE instrument (*Phylum Tech*). Each condition was tested in triplicate across three independent experimental repeats.

### ER stress vulnerability assay

NGM plates containing 1µg/mL tunicamycin (*Abcam; ab120296)* (0.0001% DMSO) were prepared (20). Synchronized L1 worms were plated and counted daily until day 1. Compound plates were compared to control NGM DMSO plates of the same genotype. The experiment was performed three times.

### Neuronal health assay

Control IZ629 *ufIs34[Punc-47p::*mCherry] were compared to mutants *srdf-3*(*mod2*)*;Punc-47p::*mCherry. Strainn were collected at day 3 of adulthood, immobilized with 5mM levamisole and visualized for motor neuron processes *in vivo* on a 2% agarose pad. Images were taken in the Cell imaging platform of the CHUM Research Center at 20X using the Zeiss *AxioObserver Z1 SDC/TIRF* and processed by *Zeiss Zen microscopy software.* Each condition was tested in triplicate (33 worms/plate) across three independent experimental repeats.

### Glycogen abundance assay

15-20 worms were exposed to iodine crystals (Sigma Aldrich; 326143) for 40-60 seconds. Images were acquired with a *Leica S6E microscope* with an iPhone 7. Staining was quantified with Fiji software by manually selecting worm images processed using the following macro: (run(“Invert”); run(“RGB Stack”); run(“8-bit”); run(“Stack to Images”); run(“Subtract Background…”, “OK”);). The experiment was performed three times.

### Liquid chromatography-tandem mass spectrometry (LC-MS/MS)

#### 1. C. elegans metabolomics

Approximately 60 mg of synchronized day 1 worms were washed three times in M9 buffer and frozen at –80 °C until extraction. Targeted metabolite analyses by LC–MS/MS were performed at The Metabolomics Innovation Center (TMIC; Alberta, Canada). Validated targeted LC–MS/MS workflows were used at TMIC to quantify mevalonate-pathway metabolites, including mevalonate, HMG-CoA, acyl-CoA intermediates, isoprenyl phosphates, and ubiquinones. Frozen worm pellets were thawed on ice and homogenized in aqueous buffer with two metal beads on an MM400 mixer mill. Aliquots of homogenate were extracted with organic solvents (e.g., methanol and methanol/dichloromethane) according to the requirements of each targeted panel. Extracts were clarified by centrifugation, and metabolite concentrations were normalized to total protein (Bradford assay). Metabolite concentrations were determined using stable-isotope internal standards and external calibration curves and quantified by multiple-reaction-monitoring LC–MS/MS. In a separate targeted assay, polyprenols and dolichols were quantified using an LC–MS/MS method optimized for long-chain polyisoprenoids. Detailed descriptions of sample preparation and LC–MS/MS parameters for all targeted panels are provided in the **Supplementary Text**. For each LC-MS/MS run, metabolite abundances were normalized to the mean value of the corresponding wild-type group, and normalized values from independent runs were pooled for analysis and visualization. For statistical comparison of metabolite abundance between wild-type and W6X worms, or between DMSO control and W6X worms, a two-sided unpaired Student’s t-test was applied, with Welch’s correction used for unequal variances. For the atorvastatin treatment experiment, mevalonate abundance was normalized to wild-type worms and analyzed by ordinary one-way ANOVA to compare atorvastatin-treated mutants to the DMSO control.

#### 2. Dolichols and polyprenols analysis from fibroblasts

Human fibroblast cell lines derived from individuals with SRD5A3-CDG and unaffected controls were obtained from previously established repositories. Written informed consent was obtained from all participants or their legally authorized representatives at the time of original sample collection, in accordance with the ethical guidelines of the originating institutions. Fibroblasts from SRD5A3-CDG patients (Supplementary Table S1) and control fibroblasts (GM5565, GM5381, GM5757, GM5400, and GM8399 (Coriell) were cultured in Minimum Essential Medium (MEM; Gibco, Carlsbad, CA, USA; 1 g/l glucose) supplemented with 10% fetal bovine serum (FBS; Gibco), 10% antibiotic-antimycotic with gentamicin, and 10% non-essential amino acids and maintained in an incubator at 37°C, 5% CO2. Cells were cultured to 80% confluence and were harvested by trypsinization with 0.05% Trypsin–EDTA (Gibco), washed with PBS and pelleted by gentle centrifugation (2000 rpm for 10 min). Dolichols and polyprenols were extracted from 1 × 10^7^ cells using the method described by Cantagrel et al (17). Cells were lysed in 50 µl water and 50 µl of 5 M KOH was added, followed by 150 µl of methanol. Cells were heated at 100°C for 60 min and 150 µl of hexane was added as extraction solvent. The organic layer was washed with water to remove any salt or protein precipitate. After washing, organic layer was dried using vacuum centrifuge and kept at -80°C until further analysis by LC-MS/MS using Orbitrap Fusion Tribrid IQ-X and Vanquish Horizon UHPLC (*Thermo Fisher Scientific*). Detailed methods on LC-MS/MS for analysis of dolichols and polyprenols from fibroblasts can be found in **Supplementary Text**. Dolichols and polyprenols were detected in the forms of ammonium adducts, [M+NH_4_]^+^, in positive ion mode electrospray. Lipid abundance was determined by calculating the peak area of the extracted ion chromatograms for the analytes at the full scan MS, using the theoretical precursor ion masses. Skyline was used for extracting and calculating the peak areas (21). The retention times were compared with those of standards to confirm the peaks. Dolichols and polyprenols containing 18-21 isoprene units were calculated. The ratios of total polyprenol/total dolichol, and individual species of polyprenol/dolichol sharing the identical number of isoprene units, were calculated. Statistical analysis was performed using a two-tailed Student t-test for comparison between unaffected control and patient fibroblasts. For statistical comparison of patient fibroblasts treated with or without atorvastatin, a paired Student t-test was applied.

### Atorvastatin supplementation

For *C. elegans*, atorvastatin (*Cedarlane; Cayman Chemicals, 20951-5*) was added to NGM plates at concentrations of 0.1, 1, 10, 50 and 100 μM. Patient fibroblasts were cultured and harvested as described above. One set was treated with 0.1 μM atorvastatin (Selleck Chemicals LLC, Cat# S5715-25mg) for 48 hours, while the other remained untreated. Scoring and imaging were blinded to genotype and/ or treatment.

### High-throughput drug screening in C. elegans

The motility-based drug screens in W6X mutant worms were conducted in 96-well plates. For the primary screen, a 3,942-compound library comprising the Prestwick Chemical Library, Sigma Aldrich LOPAC 1280 Library, Microsource Drug Library, and the BML Natural Products Library from Enzo Life Sciences was screened in singlicate. All compounds were dissolved in DMSO. ∼ 50–60 mutant adult day one worms were added to each well (1 drug/well, [20 mM], or DMSO). Swimming movement was measured every 30 minutes for 10 hours using WMicroTracker (Phylum Tech). Measurements were performed in triplicates and the average movement score was compared to the DMSO control and the average movement score of the whole plate. If the motility score for a drug was higher than the respective controls, it was selected for further validation in the secondary screen. In the secondary screen, worms were exposed to 153 compounds from the primary screen in triplicates (20 μM or DMSO). Swimming movement was measured every 30 minutes over 4 hours. Compounds that significantly increased the swimming movement in the secondary screen were considered as positive hits. Robust Z-scores were defined as: (Activity_Sample_ - Median(Activity_All within-plate samples_)) / Median absolute deviation(Activity_All within-plate samples_). For validation in the progeny assay, compounds were reordered as 25 mg dry powder stocks (Cedarlane), solubilized with fresh DMSO at 50 mM or 100 mM and stored as aliquots at −20°C. For worm retesting, a 10 mM stock was prepared.

### Data analysis and statistics

Analyses were performed in GraphPad Prism 9. Statistical significance for comparing multiple groups was assessed using one- or two-way ANOVA with Bonferroni or Tukey posthoc tests. Two-group comparisons were performed using a two-tailed Student’s t-test with Welch’s correction applied to samples with unequal variance. Paralysis and lifespan curves were analyzed using the Log-rank (Mantel-Cox) test. Data are presented as mean ± SEM.

## Data availability

Source data are provided as Supplementary Data files. Additional data are available from the corresponding author upon reasonable request.

## Acknowledgments

This study was funded by the Sappani Foundation and Cure SRD5A3. We gratefully acknowledge Claudia Maios for liquid culture experiments efforts and Audrey Labarre for technical and academic assistance. We thank Hudson Freeze and Bobby Ng for providing the fibroblast materials. We thank the *Caenorhabditis Genetics Center* (University of Minnesota), funded by NIH Office of Research Infrastructure Programs (P40 OD010440) for providing some strains.

## Author contributions

E.P., E.S., J.D., and K.K. conceived the study. A.P., J.A.P., E.M., E.P., E.S., H.D. and K.K. designed the experiments and analyses. H.D., C.M., J.D., J.P., S.K.B., and W.R. performed the experiments. H.D., E.P., K.K. and S.K.B. analyzed the data and generated the figures. H.D., KK. And S.K.B. wrote and prepared the manuscript. All authors have reviewed and approved the manuscript.

## Ethics declarations

Competing Interests: None declared.

