## Supplementary Information for "Repurposing the HMG-CoA Reductase Inhibitor Atorvastatin for SRD5A3-CDG"

***Supplementary Text***

*Liquid chromatography-tandem mass spectrometry (LC-MS/MS) for metabolites in worms*

Targeted LC–MS/MS profiling of mevalonate-pathway metabolites and polyisoprenoids was performed at The Metabolomics Innovation Center (TMIC). TMIC used targeted LC-MS/MS workflows optimized for quantification of HMG-CoA, mevalonate, acyl-CoA intermediates, isoprenyl phosphates, and ubiquinones, each employing stable-isotope internal standards and external calibration curves. A dedicated targeted assay was used for quantitation of long-chain polyprenols and dolichols. All metabolite concentrations were normalized to total protein. Metabolite measurements were acquired in two independent runs; within each run data were normalized to the corresponding wild-type controls and then pooled from downstream analysis.

*1. Quantitation of HMG-CoA and isoprenyl phosphates*

Clear supernatant was mixed 1:1:1 (v/v/v) with HMG-CoA-D3 internal standard solution and dichloromethane (DCM), vortexed for 30 s and centrifuged for 5 min. The upper aqueous phase was dried under nitrogen, then reconstituted in 50 µL of 50% methanol. Calibration standards containing HMG-CoA, acetoacetyl-CoA, acetyl-CoA, and isoprenyl phosphates were prepared by serial dilution in a ratio of 1 to 5 (v/v). Samples and calibration standards (15 µL) were analyzed by MRM-based targeted metabolomics using an Acquity UPLC (*Waters*) coupled to a QTRAP 6500 mass spectrometer (*Sciex)* operated in negative-ion mode, with a $C_{18}$ column (2.1×100 mm, 1.7 µm) and a mobile phase of 2mM ammonium acetate (A) and acetonitrile (B) under a binary gradient (2% to 60% B in 12 min) at 0.35 mL/min and 50 °C. Concentrations of the detected compounds were calculated by interpolating the constructed linear-regression, internal-calibration curves with the peak area ratios measured from sample solutions. IPP and DMAPP co-eluted; total peak area is reported.

*2. Quantitation of ubiquinones*

Supernatants were extracted twice using a mixture of 250 µL 50% methanol and 500 µL dichloromethane (DCM) (1:1:2, v/v/v). The pooled DCM layers were dried under nitrogen and reconstituted in 50 µL isopropanol. Calibration standards for MK-4, CoQ8, CoQ9, and CoQ10 mixture of dolichols 13-21 mixture of polyprenols 13-21, were prepared in isopropanol parallel by serial dilution in a ratio of 1 to 5 (v/v) as part of TMIC's standard calibration mixture. Only ubiquinones were analyzed for this study: although MK-4 and long-chain polyprenol/dolichol standards were included in the TMIC calibration mixture, MK-4 is not produced by C. elegans and polyprenols were below the detection limit in this panel. Samples and calibration standards (20 µL) were analyzed by LC-MRM/MS using an Acquity UPLC (*Waters*) coupled to a QTRAP 6500 mass spectrometer (*Sciex*) in positive-ion mode, with a $C_8$ column (2.1× 50 mm, 1.7 µm) and a mobile phase of 0.1% formic acid in water (A) and 0.1% formic acid in isopropanol (B) under a binary gradient (50% to 95% B in 15 min) at 0.25 mL/min and 50 °C. Concentrations of the detected compounds were calculated by interpolating the constructed linear-regression calibration curves from the injection of calibration solutions, with the peak areas measured from sample solutions.

*3.. Quantitation of polyprenols and dolichols*

Calibration standards were prepared by serial dilution of dolichol (C13–C21) and polyprenol (C13–C21) standards in methanol–chloroform (1:1, v/v). Frozen worm pellets were thawed, diluted (500 µL water per 50 mg tissue), and homogenized at 30 Hz for 3 min using an MM400 mixer mill**.** Two sequential hexane extractions (400 µL each) were performed; pooled organic phases were dried under nitrogen and reconstituted in 30 µL methanol–chloroform (1:1)**.** Protein in the residual aqueous pellet was measured by BCA assay for normalization. Samples and calibration standards (12 µL) were analyzed by LC–MRM/MS using an Agilent 1290 UHPLC coupled to a Triple Quadrupole 7500 mass spectrometer (*Sciex*), in positive-ion mode, a $C_8$ UPLC column (2.1× 50 mm, 1.8 µm) and a mobile phase of 2mM ammonium acetate with 0.04% acetic acid in water (A) and a 2mM ammonium acetate with 0.04% acetic acid in acetonitrile:isopropanol:water (300:700:2) (B) under a binary gradient (50% to 75% B from 0-10 min; 75% to 100% B from 10-22 min; 100% B for 1 min) at ) at 0.3 mL/min and 55 °C. The column was reconditioned at 50% B for 4.5min between injections. Peak areas were integrated. The concentrations of total dolichols and total polyprenols detected in the samples were calculated with external standard calibration.

*LC-MS/MS for analysis of dolichols and polyprenols from fibroblasts*

Dolichols and polyprenols were analyzed from unaffected control and patient fibroblasts using Vanquish Horizon UHPLC (*Thermo Fisher Scientific*) coupled to an Orbitrap Fusion Tribrid ID-X mass spectrometer (*Thermo Fisher Scientific*). A 10 min-long targeted method was developed using dolichol and polyprenol standards (*Avanti Polar Lipids*). Extracted lipids were loaded onto a Hypersil Gold Vanquish $C_{18}$ UHPLC column (2.1 × 150mm, 1.9 µm) with flow rate of 300 µl/min. Under a binary gradient of mobile phase A ($H_2O$:acetonitrile=4:6 (v/v) with 1 mM ammonium acetate) and mobile phase B (isopropanol:methanol:acetonitrile=8:1:1 (v/v/v) with 1 mM ammonium acetate), separation of dolichols and polyprenols was carried out. Mobile phase B from increased from 60 to 85% over 1 min, 85 to 100% over 4 min and maintained at 100% for 2 min. Then, the mobile phase B was decreased from 100 to 40% over 0.1 min and equilibrated at 40% for 2.9 min for the next injection. The column temperature was set at 50°C throughout the analysis. Precursor scans were acquired in the range of 500 to 1,800 m/z at 60,000 resolution (400 m/z) and targeted MS/MS scans at collision energy of 25% were acquired from the precursor m/z of dolichols and polyprenols at 15,000 resolution (400 m/z). Automatic gain control settings for MS and MS/MS were set at $4 \times 10^5$ and $5 \times 10^4$, respectively, with injection times to reach automatic gain control were set at 50 ms and 22 ms, respectively. For the analysis of dolichols and polyprenols from patient fibroblasts, both untreated and treated with atorvastatin, LC-MS/MS data acquisition was performed using an Orbitrap Fusion Tribrid IQ-X mass spectrometer (*Thermo Fisher Scientific*).

*Supplementary Figures and Tables*

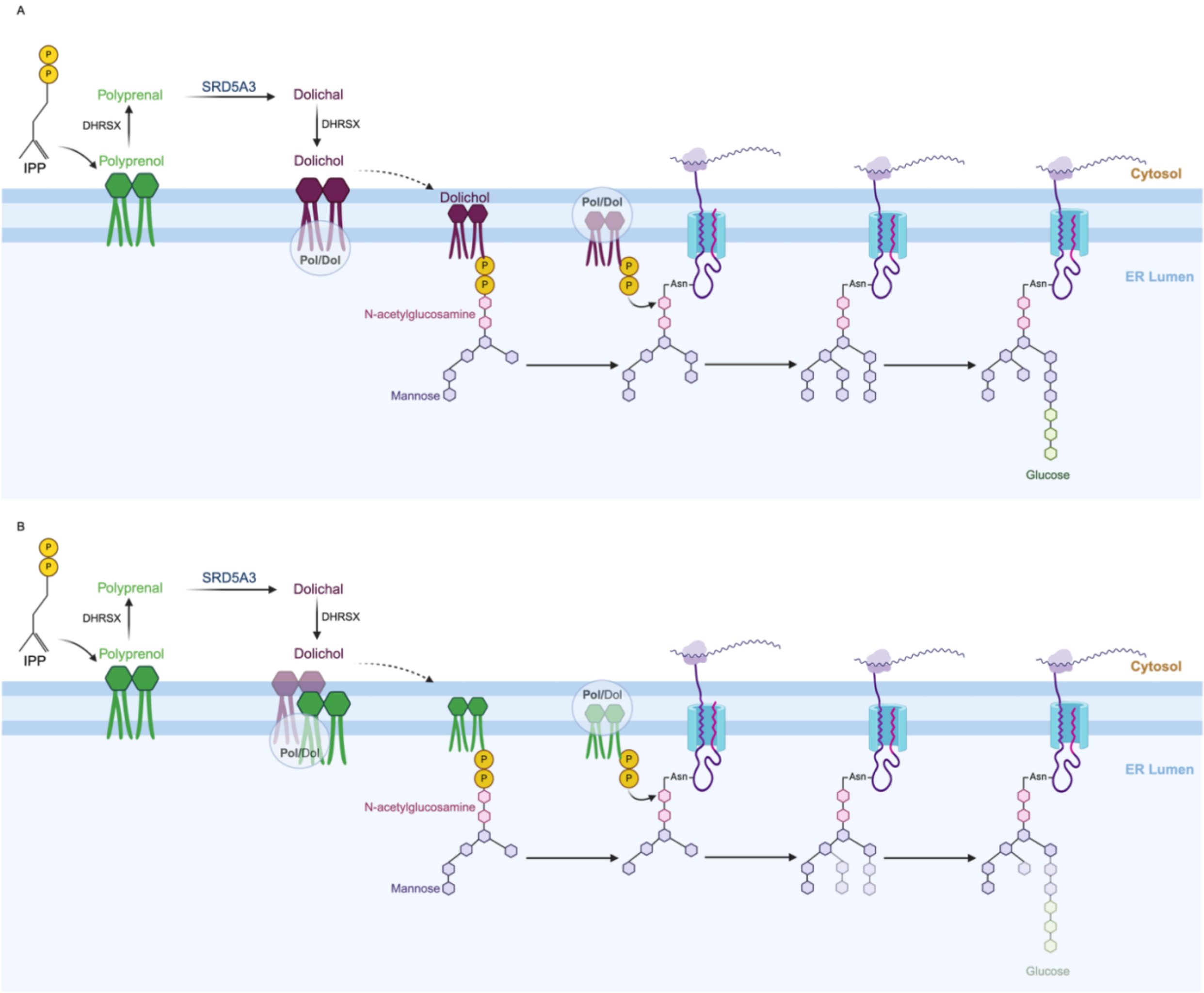


**Figure S1. Role of SRD5A3 in glycosylation and SRD5A3-CDG.** (**A**) N-glycosylation is a post-translational modification where glycans are attached to asparagine residues on proteins in the endoplasmic reticulum. Dolichol-phosphate is a lipid carrier of the oligosaccharide prior to its transfer to proteins. Dolichol is synthesized through a multi-step process: DHRSX functions as both a dehydrogenase and reductase, while SRD5A3 catalyzes the reduction of polyprenal to dolichal. (**B**) SRD5A3-CDG is caused by mutations in the SRD5A3 gene, leading to the accumulation of polyprenal and polyprenol. Phosphorylated polyprenol competes with dolichol as an oligosaccharide acceptor, resulting in inefficient glycosylation and the transfer of immature glycans to proteins. *Created with BioRender.com*

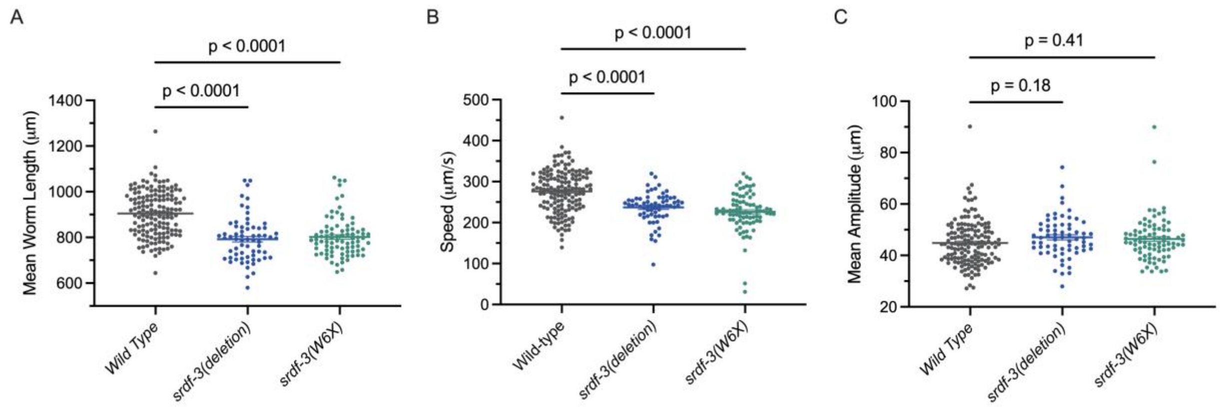


**Figure S2. Motility phenotypes of *srdf-3* mutant worms.** *srdf-3* mutant worms exhibit significant differences in mean worm length (**A**) and speed (**B**) but not in mean amplitude (**C**) relative to wild-type worms. **a-c:** $n$ ≈30 worms per experiment, $N = 3$ independent experiments. One-way ANOVA with Bonferroni *post hoc* test. Data are presented as mean ± SEM.

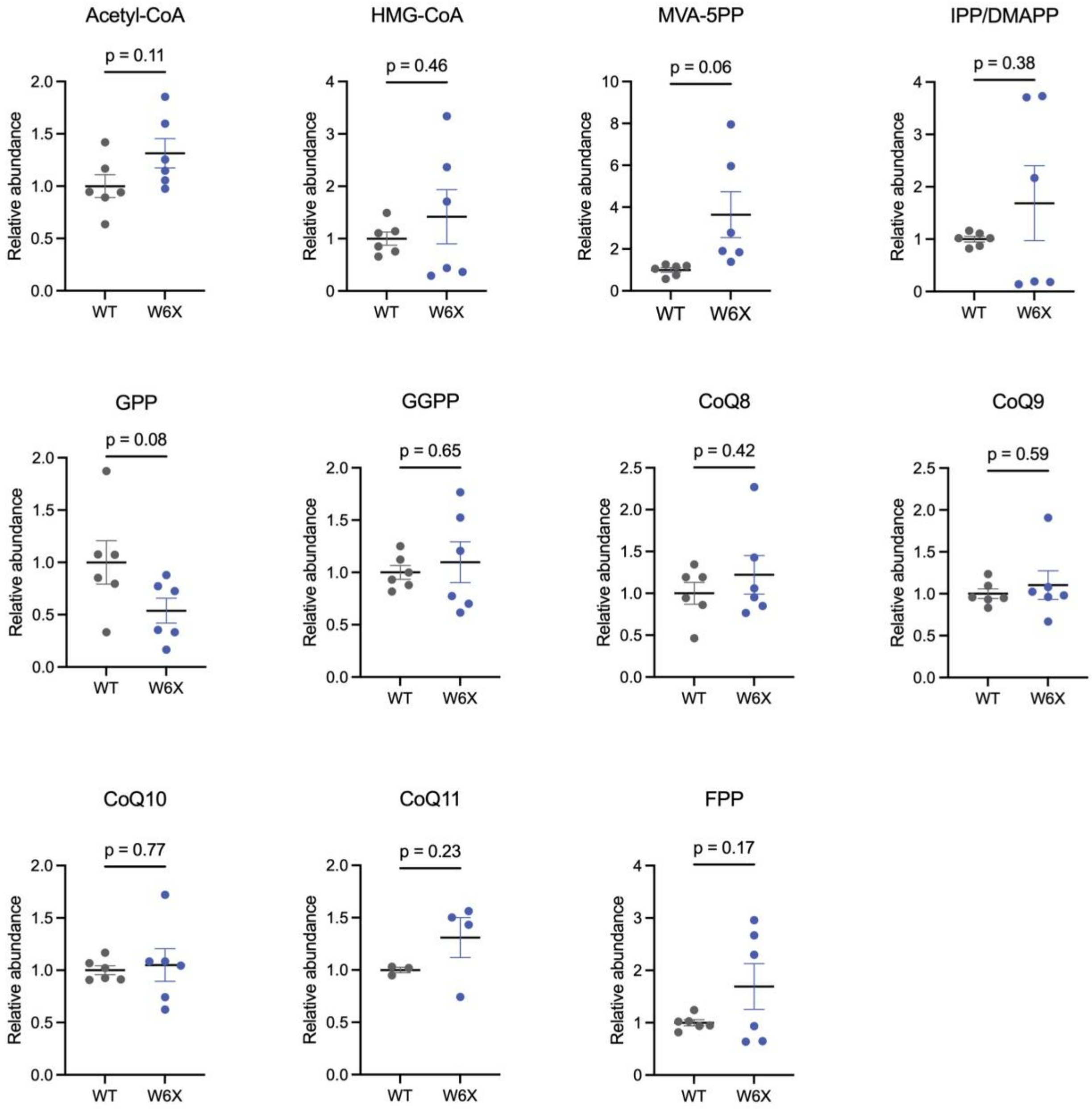


**Figure S3. Metabolomic profiling of mevalonate and dolichol biosynthesis pathway metabolites in W6X worms.** LC-MS/MS quantified metabolites in W6X worms relative to wild-type (WT) animals. $n = 3$ technical replicates, $N = 2$ independent experiments. Two-sided unpaired t-test (Welch's Correction applied where appropriate). Data are presented as mean ± SEM.

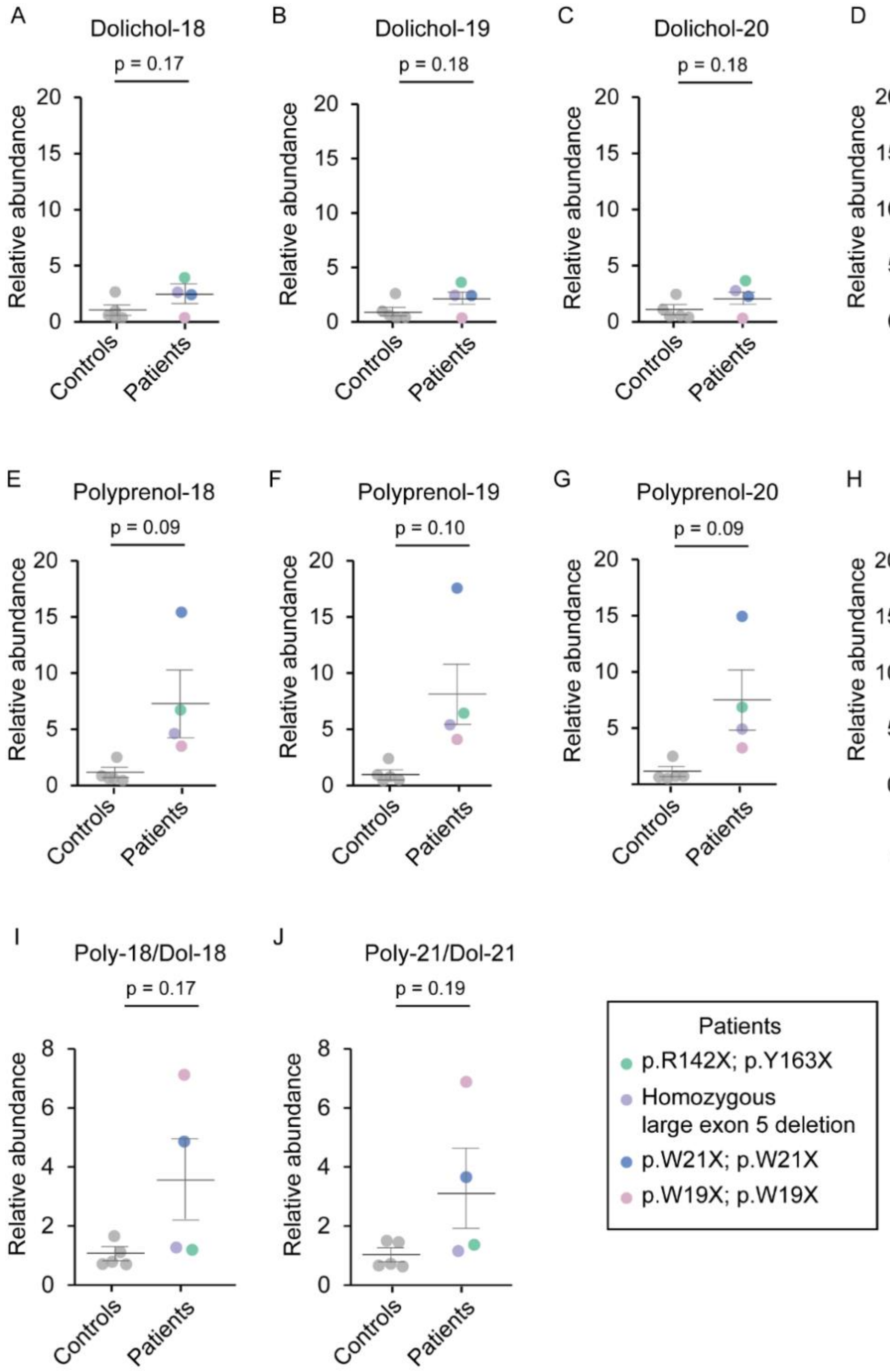


**Figure S4. Analysis of dolichols and polyprenols from SRD5A3-CDG patient and control fibroblasts.** Relative abundance of (**A**) dolichol-18, (**B**) dolichol-19, (**C**) dolichol-20, (**D**) dolichol-

21, (**E**) polyprenol-18, (**F**) polyprenol-19, (**G**) polyprenol-20, and (**H**) polyprenol-21 to unaffected controls, quantified by LC-MS/MS. Ratios of (**I**) polyprenol-18 to dolichol-18 and (**J**) polyprenol-21 to dolichol-21 are shown. Patient genotypes are indicated. **a-j**: n = 1 sample per cell line, *N* = 1 experiment. Two-sided unpaired t-test (Welch's Correction applied where appropriate). Data are presented as mean ± SEM.

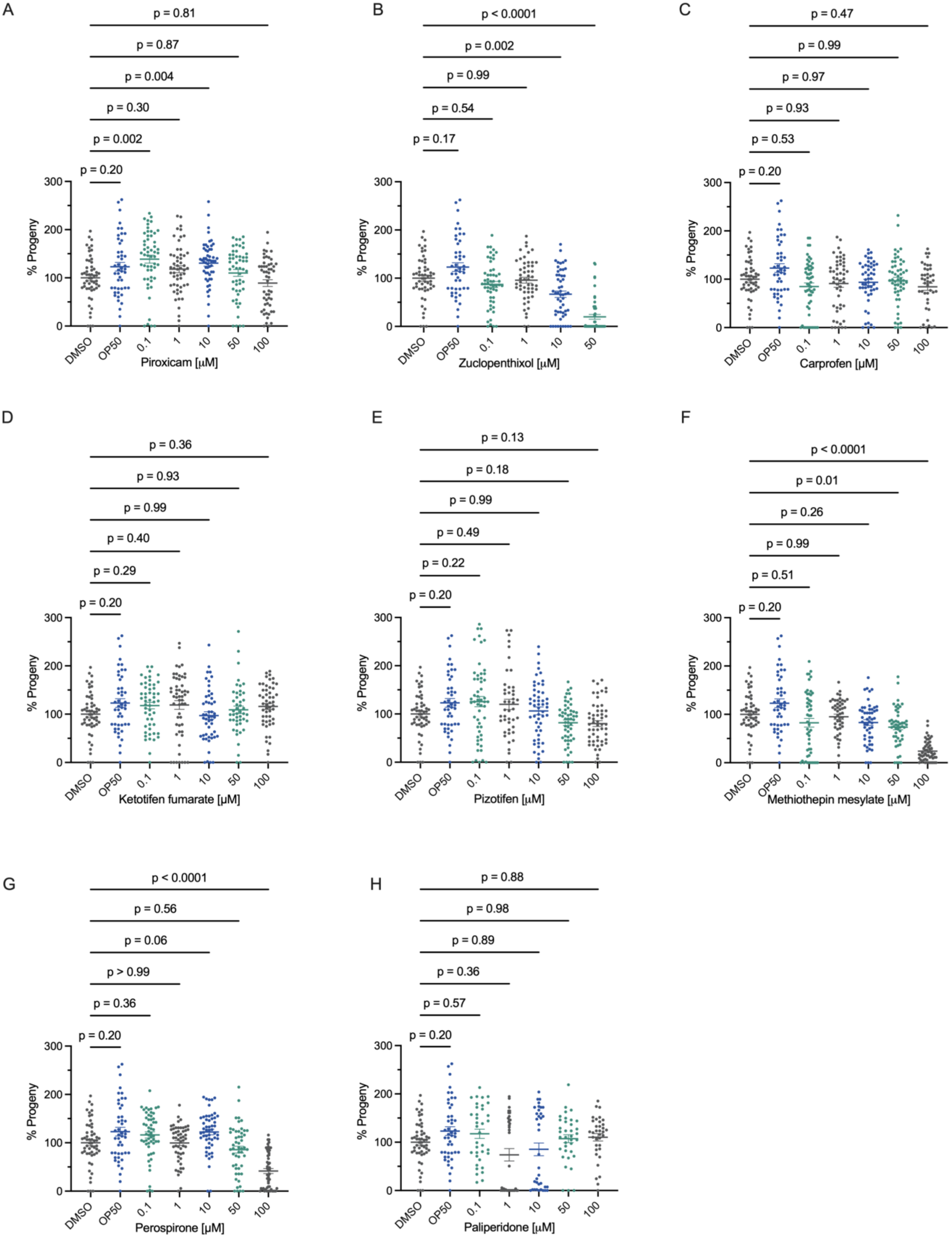
A
p = 0.81
p = 0.87
p = 0.004
p = 0.30
p = 0.002
p = 0.20
% Progeny
0
100
200
300
DMSO
OP50
0.1
1
10
50
100
Piroxicam [µM]
B
p < 0.0001
p = 0.002
p = 0.99
p = 0.54
p = 0.17
% Progeny
0
100
200
300
DMSO
OP50
0.1
1
10
50
Zuclopenthixol [µM]
C
p = 0.47
p = 0.99
p = 0.97
p = 0.93
p = 0.53
p = 0.20
% Progeny
0
100
200
300
DMSO
OP50
0.1
1
10
50
100
Carprofen [µM]
D
p = 0.36
p = 0.93
p = 0.99
p = 0.40
p = 0.29
p = 0.20
% Progeny
0
100
200
300
DMSO
OP50
0.1
1
10
50
100
Ketotifen fumarate [µM]
E
p = 0.13
p = 0.18
p = 0.99
p = 0.49
p = 0.22
p = 0.20
% Progeny
0
100
200
300
DMSO
OP50
0.1
1
10
50
100
Pizotifen [µM]
F
p < 0.0001
p = 0.01
p = 0.26
p = 0.99
p = 0.51
p = 0.20
% Progeny
0
100
200
300
DMSO
OP50
0.1
1
10
50
100
Methiothepin mesylate [µM]
G
p < 0.0001
p = 0.56
p = 0.06
p > 0.99
p = 0.36
p = 0.20
% Progeny
0
100
200
300
DMSO
OP50
0.1
1
10
50
100
Perospirone [µM]
H
p = 0.88
p = 0.98
p = 0.89
p = 0.36
p = 0.57
p = 0.20
% Progeny
0
100
200
300
DMSO
OP50
0.1
1
10
50
100
Paliperidone [µM]

**Figure S5. Dose-response of compounds on progeny numbers in W6X worms. (A)** Piroxicam treatment rescues the progeny phenotype of W6X worms at 1 μM and 10 μM doses. $n = 20$ worms per experiment, $N = 3$ independent experiments. One-way ANOVA with Bonferroni *post hoc* test. Data are presented as mean ± SEM. **(B-H)** Dose-response effect of selected compounds on the progeny phenotype. **a-h**: $n \approx 20$ worms per experiment, $N = 3$ independent experiments. Data are presented as mean ± SEM. Vehicle-control "DMSO"; Bacterial food source control "OP50".

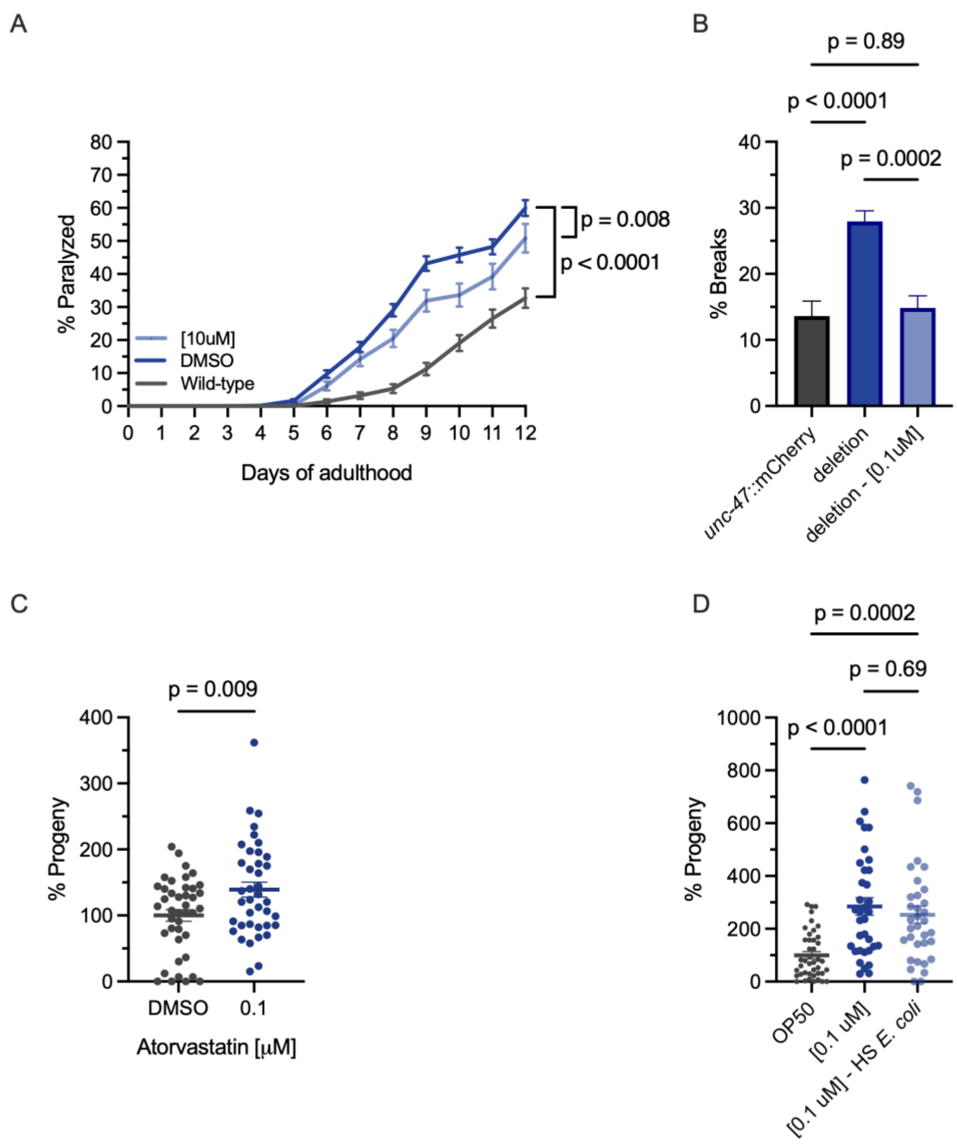


**Figure S6. Impaired movement and neuronal function in *srdf-3(deletion)* worms. (A)** Deletion mutants cultured on solid media show motility defects leading to paralysis at a higher rate compared to wild-type controls. $n \approx 120$ worms per experiment, $N = 3$ independent experiments. Atorvastatin (10 µM) rescues paralysis in deletion mutant worms. Log-rank (Mantel-Cox) test two-sided. Data are presented as mean ± SEM. **(B)** Deletion mutants have a greater frequency of breaks/gaps along neuronal processes compared to wild-type *unc-47::*mCherry controls at day 3 of adulthood which is rescued with atorvastatin treatment 0.1 µM). $n \approx 30$ worms per group, $N = 3$ independent experiments. Brown-Forsythe one-way ANOVA with Dunnett *post hoc* test. Data are presented as mean ± SEM. **(C)** Atorvastatin (0.1 μM) rescues the progeny phenotype in deletion mutant worms. $n \approx 20$ worms per experiment, $N = 3$ independent experiments. Two-sided unpaired t-test. Data are presented as mean ± SEM. **(D)** Heat-shock killed *E. coli* culture does not impact the rescue effect of atorvastatin (0.1 μM) treatment on the progeny phenotype in deletion mutants. $n \approx 20$ worms per experiment, $N = 3$ independent experiments. Brown-Forsythe one-way ANOVA with Tukey *post hoc* test. Data are presented as mean ± SEM. Vehicle-control "DMSO"; Bacterial food source control "OP50".

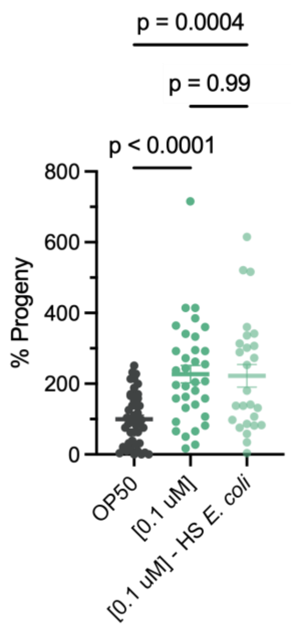


**Figure S7. Rescue of the progeny phenotype in W6X mutant worms by atorvastatin is independent of *E. coli.*** Heat-shock killed *E. coli* culture does not impact the rescue effect of atorvastatin (0.1 μM) treatment on the progeny phenotype in W6X mutants. $n \approx 20$ worms per experiment; $N = 3$ independent experiments. One-way ANOVA with Tukey *post hoc* test. Data are presented as mean ± SEM. Bacterial food source control "OP50".

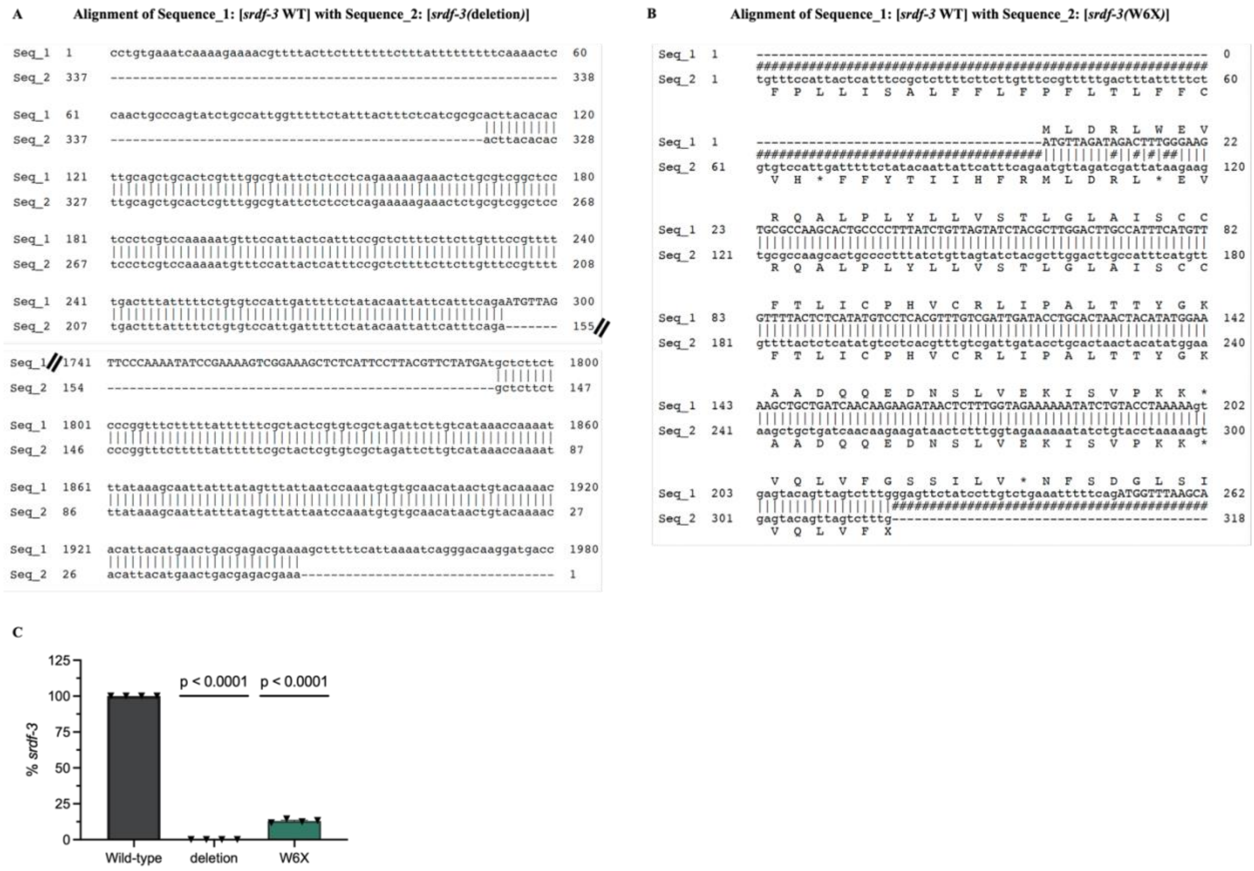


**Figure S8. Genotyping evidence of *srdf-3* mutants. (A)** Genotyping and sequence alignment of *srdf-3(deletion)* and **(B)** srdf-3(W6X) mutants show the localization of each mutation. **(C)** Quantitative PCR shows a significant decrease in mRNA levels of *srdf-3* in the deletion and W6X mutants. $N = 4$ independent experiments. Two-sided one-sample t-test. Data are presented as mean ± SEM.

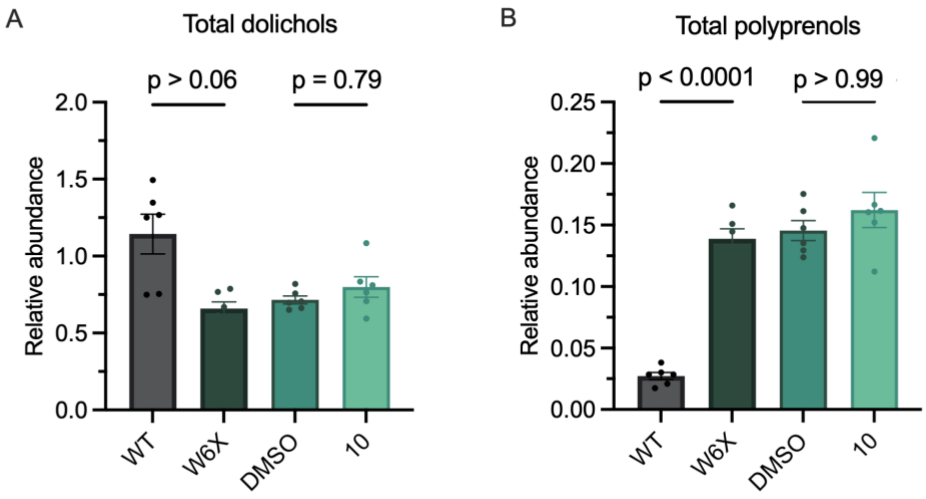


**Figure S9. Atorvastatin does not alter dolichol or polyprenol abundance in W6X worms.** Atorvastatin (10μM) does not significantly change **(A)** total dolichols levels or **(B)** total polyprenols levels in W6X mutant worms relative to DMSO-treated control, quantified by LC-MS/MS. **a-b**: *n* = 3 technical replicates, *N* = 2 independent experiments. Brown-Forsythe one-way ANOVA with Dunnett *post hoc* test. Data are presented as mean ± SEM. Vehicle-control "DMSO"; wild-type "WT".

**Table S1. Genetic and clinical details of SRD5A3-CDG patients.** List of individuals with SRD5A3-CDG used in the study along with genetic information and clinical characteristics.

| | Case 1 | Case 2 | Case 3 | Case 4 |
|---|---|---|---|---|
| **Age (y)** | 4 | 7 | 9 | 17 |
| **Sex** | F | F | M | M |
| **Predicted protein variant in SRD5A3** | Homozygous large deletion in Exon 5 | p.W21X; p.W21X | p.R142X; p.Y163X | p. W19X; p. W19X |
| **Clinical characteristics** | | | | |
| Elevation in AST/ALT | + | + | + | + |
| Abnormal Coagulation | + | + | + | + |
| Cerebellar hypoplasia | + | + | + | + |
| Coloboma and visual loss | + | + | + | + |
| Developmental delay | + | + | + | + |
| Intellectual disability | + | + | + | + |
| Ichthyosis | + | - | + | - |
| Microcytic anemia | + | - | - | - |

*Abbreviations: AST – aspartate transaminase; ALT - alanine transaminase

**Table S2. Z-score and average movement activity across all timepoints of selected compounds identified as positive hits in the secondary screen.**

| Compound | Z-Score | Activity (% of negative control) |
|---|---|---|
| Methiothepin mesylate | 2.7856 | 150.00 |
| Thioridazine hydrochloride | 2.2033 | 130.23 |
| Irigenin trimethyl ether | 2.0235 | 108.86 |
| Pizotifen malate | 1.4416 | 128.97 |
| Perospirone | 1.3490 | 124.86 |
| Zuclopenthixol dihydrochloride | 1.2829 | 127.78 |
| Chlorpromazine hydrochloride | 1.2141 | 124.01 |
| Syringetin-3-glucoside | 1.1768 | 112.35 |
| Cyproheptadine hydrochloride | 1.0687 | 129.84 |
| Carprofen | 0.9390 | 125.20 |
| Hypocrellin A | 0.8898 | 110.36 |
| Loxapine succinate | 0.8409 | 127.16 |
| Piroxicam | 0.7533 | 126.13 |
| Atorvastatin | 0.6745 | 120.62 |
| Hypocrellin B | 0.6745 | 118.90 |
| Chlorprothixene hydrochloride | 0.5951 | 122.62 |
| Isotectorigenin, 7-methyl ether | 0.4497 | 100.59 |
| Ketotifen fumarate | 0.4205 | 122.22 |
| Paliperidone | 0.3504 | 121.40 |
| Loratadine | 0.1984 | 119.64 |
| Acetylsalicylic acid | 0.1719 | 119.44 |
| Dicourmarin | 0.1148 | 104.98 |

**Table S3.** ***C. elegans*** **strains and genotypes**

| Strain | Genotype | Available from |
|---|---|---|
| N2 | Wild type *C. elegans* strain | CGC |
| MOD1 | *srdf-3(mod1)* | Parker Lab |
| MOD2 | *srdf-3(mod2)* | Parker Lab |
| MOD3 | *srdf-3*(*mod1*)*;Punc-47p::*mCherry | Parker Lab |
| MOD4 | *srdf-3*(*mod2*)*;Punc-47p::*mCherry | Parker Lab |
| IZ629 | *ufIs34[Punc-47p::*mCherry] | Kind gift from Dr. Michael M. Francis (University of Massachusetts, Worcester, MA) |
| CB246 | *unc-64(e246)* | CGC |
| CB307 | *unc-47(e307)* | CGC |
| CB1072 | *unc-29(e1072)* | CGC |
